# Type-1 IFN primed monocytes in pathogenesis of idiopathic pulmonary fibrosis

**DOI:** 10.1101/2020.01.16.908749

**Authors:** Emily Fraser, Laura Denney, Karl Blirando, Chaitanya Vuppusetty, Agne Antanaviciute, Yuejuan Zheng, Emmanouela Repapi, Valentina Iotchkova, Stephen Taylor, Neil Ashley, Victoria St Noble, Rachel Benamore, Rachel Hoyles, Colin Clelland, Joseph M D Rastrick, Clare S Hardman, Nasullah K Alham, Rachel E Rigby, Jan Rehwinkel, Ling-Pei Ho

## Abstract

Idiopathic pulmonary fibrosis (IPF) is the most severe form of lung fibrosis. It is progressive, and has an extremely poor outcome and limited treatment options. The disease exclusively affects the lungs, and thus less attention has been focused on blood-borne immune cells. which could be a more effective therapeutic target than lung-based cells. Here, we questioned if circulating monocytes, which has been shown to be increased in IPF, bore abnormalities that might contribute to its pathogenesis. We found that levels of circulating monocytes correlated directly with the extent of fibrosis in the lungs, and increased further during acute clinical deterioration. Monocytes in IPF were phenotypically distinct, displaying increased expression of CD64, a type 1 IFN gene expression signature and a greater magnitude of type 1 IFN response when stimulated. These abnormalities were accompanied by markedly raised CSF-1 levels in the serum, prolonged survival of monocytes *ex vivo*, and increased numbers of monocytes in lung tissue. Our study defines the key monocytic abnormalities in IPF, proposing type 1 IFN-primed monocytes as a potential driver of an aberrant repair response and fibrosis. It provides a rationale for targeting monocytes and identifies monocytic CD64 as a potential specific therapeutic target for IPF.

## INTRODUCTION

Fibroproliferative diseases, including the various fibrotic lung diseases, systemic sclerosis, liver cirrhosis and macular degeneration, are leading causes of morbidity and mortality. Idiopathic pulmonary fibrosis (IPF) is the most severe form of chronic fibrotic lung disease (1). Unlike some of the other fibrotic diseases, IPF, is an unremitting, progressive disease, with a median survival of only five years from diagnosis (2, 3). It is the outcome of a complex interaction of genetic and environmental factors, ageing-associated processes and epigenetic reprogramming. Repeated but minor insults to the alveolar epithelium, for example by gastro-oesophageal reflux, viral infection or cigarette smoking, are thought to lead to a disproportionate repair response by fibroblasts and mesenchymal cells (4). This aberrant wound healing response results in parenchymal damage, an expanded collagen-rich interstitium and alveolar destruction which ultimately leads to impaired gas exchange. Research has focused more intensely on the aberrant repair response, and less on the causes of injury and drivers of chronic fibrosis. This has led to more than 40 clinical trials, but only two drugs (Nintedanib and Pirfenidone), which slow down but do not halt progression of disease (1). A better understanding of the initiators and immune drivers of fibrosis in IPF will increase the range of new therapeutic targets for this condition.

There is ample evidence that tissue macrophages are involved in fibrogenesis (5–7). Abnormalities in macrophages can lead to aberrant repair due to anomalous cross-talk with stromal cells (including epithelial cells, endothelial cells, fibroblasts, and progenitor cells), with exuberant production of inflammatory mediators and growth factors (6). Recent single cell technology has also indicated that there are specific pro-fibrotic macrophage subsets (in murine models), which drive fibroblast proliferation by secreting platelet-derived growth factor (PDGF) (8). Unsurprisingly, lung macrophages have been proposed as a therapeutic target in IPF (9). However, in practice, these lung-based cells, some, in dense matrix-rich interstitium (10) may be poorly accessible to potential drugs. In contrast, a cellular target in the blood, show to be linked to fibrosis in the lungs could be an ideal drug focus. In this regard, the precursor of lung macrophages – the circulating monocyte, is of immense interest.

Monocytes play important roles in immune defense, inflammation and homeostasis (11). Typically, they are quickly mobilized to sites of injury or infection where they contribute to the initial inflammatory processes and instigate adaptive immune responses. Once in tissues, they rapdily differentiate into macrophages in response to local environmental cues (12), producing macrophages with specific functions, e.g. macrophages that facilitate healing or those that restore tissue homeostasis after lung injury (13–15). Monocyte-derived, rather than embryonically-derived alveolar macrophages, have now been clearly shown to be critical for the development of lung fibrosis (in mice), and potentially also in the resolution of fibrosis (5, 16)

Relatively little is known about the contribution of monocytes *per se* (rather than macrophages) to lung fibrosis in human disease. These immune cells are largely inflammatory and injurious though can also aid repair of damaged tissue. In organs other than lungs, animal studies have described the ability of monocytes to enhance myofibroblast proliferation in cardiac muscles after infarction (17), and to promote progenitor regrowth after eye and spinal cord injury mediated via production of IL-10 (18). In recent years, it has also become clear that during conditions of immune ‘stress’ e.g. severe inflammation or cancer, the generation of monocytes in the bone marrow can bypass canonical routes to produce immature forms with altered function, some with pro-fibrotic features (19). Thus, monocytes in their own right, could be important effectors in fibrosis, both by causing or enhancing injury, and thus driving repair, and also by differentiating into pro-fibrotic macrophages. A recent study has shown increased levels of circulating monocytes in IPF patients, which was associated with poorer survival, but not a potential mechanism for how monocytes might contribute to pathogenesis, nor a direct link to fibrosis (42).

In this study, we investigate whether monocytes in IPF patients bear abnormalities that might contribute to the pathogenesis of IPF. We found that circulating monocytes levels correlated directly with the extent of fibrosis in the lungs. In addition, they are phenotypically distinct, displaying increased expression of CD64, a type 1 IFN signature in their transcriptome and a greater magnitude of type 1 IFN response when stimulated. These abnormalities are accompanied by markedly raised CSF-1 levels in the serum and prolonged survival of monocytes *ex vivo*. Our study defines the key monocytic abnormalities in IPF, proposing type 1 IFN-primed monocytes as possible mediators of injury, and a potential driver of an aberrant repair response and fibrosis. It provides a rationale for monocytes as a therapeutic target in IPF.

## RESULTS

### Circulating monocytes levels in IPF correlate with the extent of lung fibrosis and display increased expression of CD64

Patients with rigorously defined IPF (see Supplemental Methods), were recruited from the Oxford Interstitial Lung Disease Clinical Service (n=37). Current smokers (or ex-smokers within 5 years), those on immunosuppressants or corticosteroids, and those with concomitant lung disease or immune conditions or cancers were excluded (demographics in Supplemental Table 1). 14 patients were on the anti-fibrotic drug (Pirfenidone); no patients were on immunosuppressants or corticosteroids. Data were analysed for all IPF patients (n=37) vs age-matched healthy controls (HC) (n=28), and also after dividing patients into those on Pirfenidone treatment (n=14) compared to those without (n=14).

Peripheral blood mononuclear cells (PBMCs) were isolated by Ficoll separation and analysed immediately without storage. Monocytes were defined as CD14^lo-hi^ CD16^neg-hi^ cells (Figure 1A). In support of Scott’s findings (42), total monocytes levels were higher in IPF patients compared to age-matched heathy controls (HC); both when quantified as % of PBMCs (Figure 1A) and absolute number (per ml of blood) (Figure 1B) [16%(5) v 12%(6); IPF vs HC; p=0.022 and 3.2×10^5 (1.6) v 1.7×10^5 (0.7)/ml; p<0.001 respectively; mean(S.D.)]. There was no difference comparing patients on Pirfenidone to those who were not (Figure S1a,b).

**Fig. 1.**
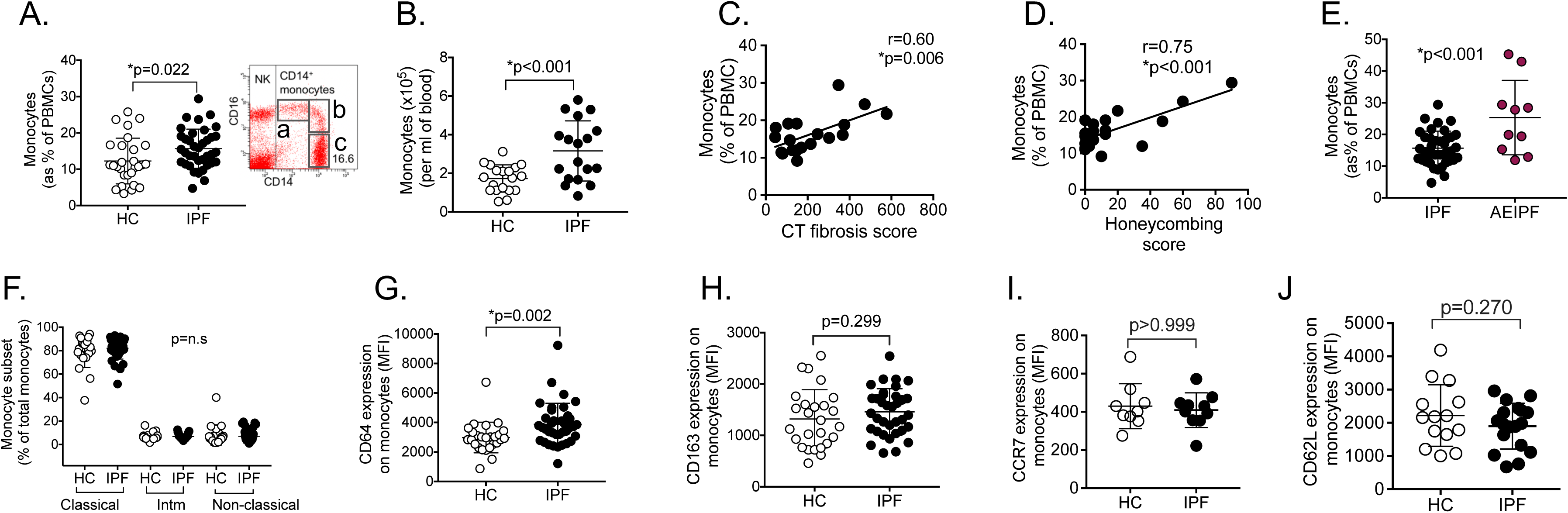
IPF monocytes are increased and correlate with extent of fibrosis and severity. (A) Monocytes levels in IPF (n=37) vs healthy controls/HC (n=28) as % of PBMC and gating for monocytes. FACS plot was gated on all live PBMCs; all monocytes analysed immediately without storage; ‘a’-non-classical monocytes, ‘b’ – intermediate monocytes, ‘c’ – classical monocytes (B) Monocytes as absolute number in blood (n=18). (C)-(D) Monocyte levels correlated positively with amount of fibrosis on contemporaneous thoracic CT scans, as measured by a fibrosis score incorporating all CT features of fibrosis (C) and also when only honeycombing fibrotic feature is quantified (n=19). (E) Increased levels of monocytes in patients with AEIPF (acute exacerbation of IPF) (n=10) compared to stable/non-AEIPF patients(IPF) (n=28). (F) Monocyte subsets in IPF and HC, showing no significant difference for any of the subsets. ‘Intm’ – intermediate. Populations shown in (A). (G)-(J) Expression of CD64 and CD163 (n=37) and CCR7 and CD62L on monocytes from a subset of IPF patients (n=9 and 18 respectively), determined by flow cytometry. p values derived using Student t test for normally distributed data, or Mann Whitney Rank Sum test otherwise. Correlation analysed using Pearson correlation test.

To examine the correlation between monocyte frequency and the extent of lung fibrosis we used a scoring method to quantify the amount of fibrosis in IPF lungs in a subset of patients (n=19). The score comprised the sum extent of reticulation, honeycombing and traction bronchiectasis [archetypal features of usual interstitial pneumonia (UIP) pattern fibrosis in IPF lungs] on high resolution thoracic computer tomographic (HRCT) scans (see Supplemental Methods). We found that the extent of fibrosis in the patients’ lungs correlated positively with levels of circulating monocytes (Figure 1C). There was an even stronger correlation between monocyte levels and total amount of honeycombing on HRCT scans (Figure 1D). Although not statistically significant, there was also a trend between higher monocyte levels and worse lung function - forced vital capacity (FVC) and diffusion capacity (TLCO) (Figure 1E-F). As TLCO and FVC can be affected by factors other than the amount of fibrosis (e.g. cardiac failure, obesity and emphysema), these results suggest that monocyte level in specifically linked to the extent of lung fibrosis.

During the study period, we were also able to recruit 10 patients who were hospitalised with acute exacerbation of IPF (AEIPF), a distinct and severe episode of acute clinical deterioration, characterised by a picture of massive injury on CT scan (20) (see Supplemental Methods). These patients had evidence of typical changes (ground glass opacification) on CT scan and no evidence of infection by aerobic and anaerobic blood cultures. Monocyte levels were measured in these patients and found to be significantly higher than in stable IPF patients [25%(12) v 16%(5); IPF vs AEIPF; p<0.001] (Figure1G and Supplemental Table 2A,B).

We next immunophenotyped the circulating monocytes, starting with the classical division of monocytes to classical, non-classical and intermediate subtypes (11). We found no difference in the proportion of intermediate (CD14^hi^CD16^hi^), non-classical (CD14^mid^CD16^hi^) and classical (CD14^hi^ CD16 ^neg/mid^) subsets (Figure 1I) between IPF and HC. Further flow cytometry analysis focusing on phenotypic markers which might confer greater inflammatory (M1-like - CD64, CD62L, CCR7) or pro-repair (M2-like – CD163) features (21) showed no abnormality apart from an increase in CD64 expression (Figure 1H-L, Supplemental Fig 1C). To complement these flow cytometry studies, gene expression (by qPCR) of M1, M2, M2a, and M2c genes [Supplementary Table 3 (21)] was also examined in freshly isolated monocytes from IPF (n=7) vs age-matched healthy controls (n=8) (demographics and RIN of RNA in Supplemental Table 4). We observed no difference in expression in these genes compared to controls - [*TGFB1, MRC1 (CD206), CD200R1, TGM2, CD163, IDO1, TNFA, IL6, CXCL10, CCR2 and IL1R2*] (Supplemental Figure 1D) (21).

In summary, we show for the first time, in a carefully conducted study on fresh samples from patients not on immunosuppressants or corticosteroids, that monocytes levels correlated positively with extent of fibrosis in the lungs of IPF patients. Against a wide immunophenotyping screen, we observed an isolated increase in expression of CD64, a high-affinity IgG receptor (FcγRI) that triggers monocyte activation upon receptor aggregation. Overexpression of CD64 on monocytes has been linked to both inflammatory markers and disease severity in rheumatoid arthritis (22, 23) and systemic lupus erythematosus (24). To date, it has not been associated with fibroproliferative diseases such as IPF.

### Serum analysis in IPF patients shows changes that specifically reflect monocyte activity and promote monocyte generation and survival

To explore possible causes for, and consequence of, increased monocyte levels and raised monocytic CD64 expression, we examined key inflammatory and monocyte-associated factors (by multi-plex bead array) in serum from IPF patients (n=24) and healthy controls HCs (n=11) (demographic in Supplemental Table 5; 8 patients were on the antifibrotic, Pirfenidone). A set of 13 cytokines and chemokines were selected to reflect the following functional groups-(i) monocyte activation/differentiation/trafficking (CCL-2 [MCP-1], CSF-1 [M-CSF], TNF-α, IL-6, CXCL-10[IP-10]); (ii) T cell activation/pro-inflammatory/trafficking (IL-13, TNF-α, CCL-20 [MIP3A], CXCL-9[MIG], IFN-γ, IL-6, IL-1β and IFN-β), (iii) monocyte/T cell anti-inflammatory (IL-10) and (iv) granulocyte differentiation/trafficking (CSF2 [GM-CSF], CXCL8 [IL-8]).

We found a significant increase in the levels of CSF-1, IL-6, and CCL2, in IPF serum compared to healthy controls [793.7(228.5) vs 250.6(102.3) pg/ml, p<0.001;7.0(3.5-25.5) vs 1.6(1.0-3.2) pg/ml, p=0.02; and 1087.0(916.2-1186.0) vs 650.8 (531.0-770.0) pg/ml,p=0.001 respectively) (Figure 2A-C). No other analytes were significantly different from controls (Figure 2D-J). IL-13, IL-10, IL-1β and IFN-β were below detection limits for the majority of patients and controls in undiluted serum samples (Supplemental Figure 2 A-D), and were not included in further correlation studies. The outcome of all analyses was the same when patients with Pirfenidone were excluded from the analyses (Supplemental Figure 2E).

**Fig. 2.**
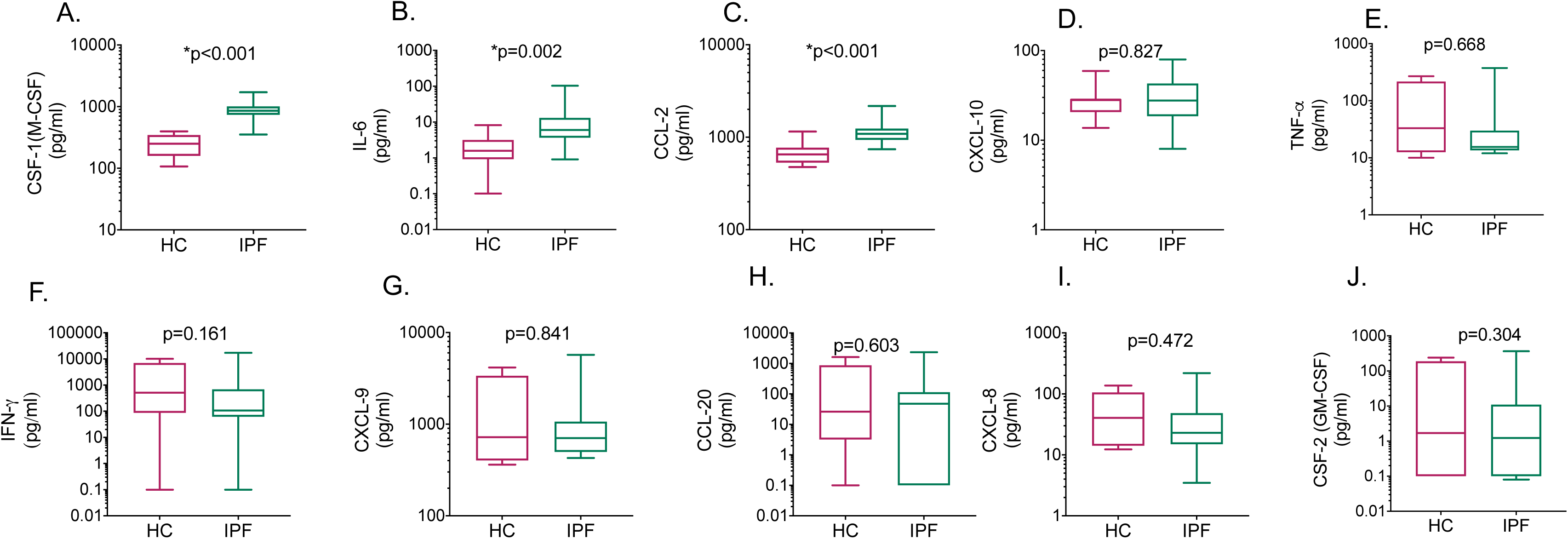
Serum profile of IPF patients show changes that reflect monocyte activity and promote monocyte generation and survival. (A)-(K) Levels of mediators measured in serum using Luminex technology showing isolated increase in factors that promote monocyte generation (CSF-1) and reflect monocytic activity (CCL-2 and IL-6) in IPF patients (n=24 IPF; n=11 healthy controls, HC) (A-C). Y axis is log 10. Zero values for serum mediators were converted to 0.1 (dotted line) for visualisation purposes on a log 10 axis. Box plot is median+/1 interquartile confidence interval, whiskers show minimum and maximum values. P values calculated using Mann Whitney Rank Sum test.

We did not observe any significant correlation between monocyte levels and any soluble mediators, nor between monocytic CD64 expression and any soluble mediators, apart from a modest but significant positive correlation between CSF-1 levels and monocytic CD64 expression in IPF patients (Supplemental Figure 2F-G).

These results show that the blood compartment in IPF patients demonstrated clear anomalies in monocyte-associated but not in lymphocytic and granulocyte-related immune mediators. Very high levels of CCL2 and IL-6 may be products of activated monocytes (25–28) while the markedly increased CSF-1 levels point to an environment that is pro-monocyte generation and survival (29).

### RNA sequencing reveals a Type 1 IFN signature in monocytes from IPF patients

In order to further investigate the abnormal monocyte phenotype detected in IPF patients, we subjected monocytes from IPF and age-matched healthy controls to bulk RNA sequencing. Monocytes were isolated by CD14^+^ magnetic bead selection (98-99% purity) from three well-characterised IPF patients (with definite UIP pattern fibrosis on CT scan) who were not on anti-fibrotics, and were non-smokers (all male; aged 57,76 and 78y) and three healthy controls (HC) (non-smokers, no medications; aged 65,68 and 71y; all males).

We focused on identifying the processes that underlie the transcriptomic changes between IPF patients and controls, rather than individual differentially expressed genes. We first subjected the differentially expressed gene list to enrichment analysis, using gene sets from the REACTOME platform, a database of signalling and metabolic molecules, organized into biological pathways and processes, with a focus on reactions. This revealed that the most significantly enriched gene sets were ‘IFN signalling’ and ‘IFNαβ signalling’ (Figure 3A and Supplemental Table 6). We tested this further by using Gene Set Enrichment Analysis (GSEA) (30) which analysed enrichment of all genes arranged according to fold change, compared to the control group, regardless of statistical significance of their differential expression. This approach also identified type 1 IFN signalling gene sets as the most enriched gene sets in IPF, using both REACTOME and GO gene sets (Figure 3B and Supplemental Table 7).

**Fig. 3.**
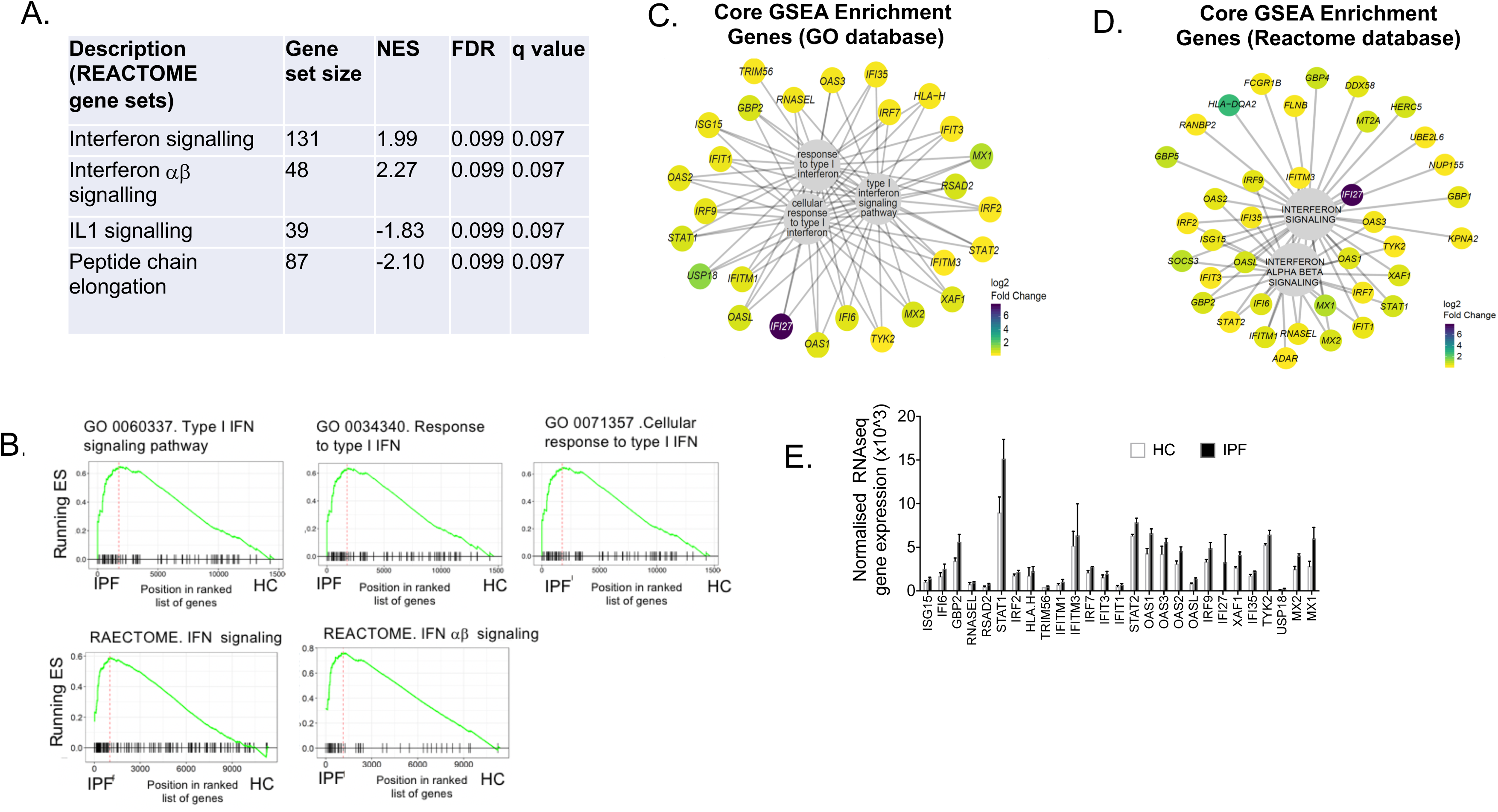
RNA sequencing of IPF monocytes reveals a Type I IFN signalling signature. (A) Analysis of differentially expressed gene list for enrichment of biologically-relevant pathway and processes using REACTOME gene sets identifies interferon signalling and IFN ab signalling as top gene sets. NES – normalised enrichment score (B) GSEA plots showing most enriched gene sets from GO and REACTOME database. ES-Enrichment score (C-D) Composition and expression levels of genes in the leading edge of the five gene sets from (B); comparing expression in IPF monocytes to control; expressed as log2 fold change. (E) Individual expression of archetypal interferon stimulated genes (ISG) from RNA sequencing data, showing increased expression in IPF compared to HC [16 archetypal ISGs (from (Sarasin-Filipowicz, Oakeley et al. 2008) and (Schoggins, Wilson et al. 2011)) and six ISGs found increased in PBMCs from patients with Aicardi-Goutières syndrome, often referred to as the type 1 IFN signature (*IFI27, ISG15, IFI44L, IFIT1, SIGLEC1* and *RSAD2*) (Rice, Forte et al. 2013].Normalisation done according to Anders (2010), using library size factors.

To explore the composition of the most highly enriched gene sets, we examined the genes that made up the core enrichment genes (i.e. the genes that contribute most to the enrichment result) in the five IFN signalling gene sets shown in Figure 3B. From these five gene sets, we observed that most of the genes were interferon stimulated genes (ISGs) from different points in the type 1 IFN signalling pathway (Figure 3D, E and Supplemental Figure 3) (31, 32). Indeed, when we examined the expression of a curated gene set of 16 archetypal ISGs [from (33) and (34)] and six ISGs found increased in PBMCs from patients with Aicardi-Goutières syndrome, [often referred to as the type 1 IFN signature (*IFI27, ISG15, IFI44L, IFIT1, SIGLEC1* and *RSAD2*)] (35), all these ISGs were also increased in IPF patients compared to HC (Figure 3F).

Thus, RNA sequencing of IPF monocytes showed that their gene expression profiles were endowed with a type 1 IFN signature. This has not been hitherto reported and is highly relevant to the observation of increased CD64 expression as type 1 IFNs can induce CD64 expression on monocytes (36), and CD64 is a biomarker of disease activity in type 1 interferonopathies like SLE (36, 37)

### Monocytes from IPF patients show greater ISG induction in response to type 1 IFN stimulation

The RNA sequencing findings suggest that monocytes from IPF patients have a primed type I IFN signalling pathway. To test this, we reasoned that IPF monocytes would respond more readily or greatly to a type I IFN stimulus. We recruited a larger cohort of IPF patients (n=27) and age-matched healthy controls (n=10), recruited on the same basis as described above (Supplemental Methods; demographics in Supplemental Table 8A). We examined gene expression by qPCR, of the most differentially expressed ISGs from our RNA sequencing data – *IFI27, USP18, MX1* and *OASL*; archetypal ISGs (*IRF7 and MX2*) and four ISGs found highly expressed in PBMCs from the type 1 interferonopathy Aicardi-Goutières syndrome (AGS) patients - *IFI27, ISG15, IFI44L and RSAD2* (38). In addition, *IFNB1 and STAT1*, a key transcription factor in the type I IFN signalling pathway were also examined.

We found that all the ISGs, *STAT1* and *IFNB1* showed higher expression in IPF compared to control, consistent with the RNA sequencing findings, with some statistically significantly increased - MX2, ISG15 and OASL (p=0.021, p=0.048 and p=0.048 respectively) (Figure 4A-K). There was no difference in findings between patients on anti-fibrotics (Pirfenidone and Nintedanib) and those not (Supplemental Figure 4A-K). As expected, expression levels of ISGs in each patient’s monocytes correlated strongly with each other, i.e those patients with high *MX1*, also had high *IRF7* and all other tested ISGs (Supplemental Figure 4T). Basal ISG expression levels were also significantly correlated with basal *IFNB1* and *IFNAR1* expression (Supplemental Figure 4T). These results provide support for the integrity of the data as these genes are expected to change contemporaneously. It is noted that neither *IFNB1* nor *IFNAR1* expression was significantly increased in IPF monocytes at homeostasis.

**Fig. 4.**
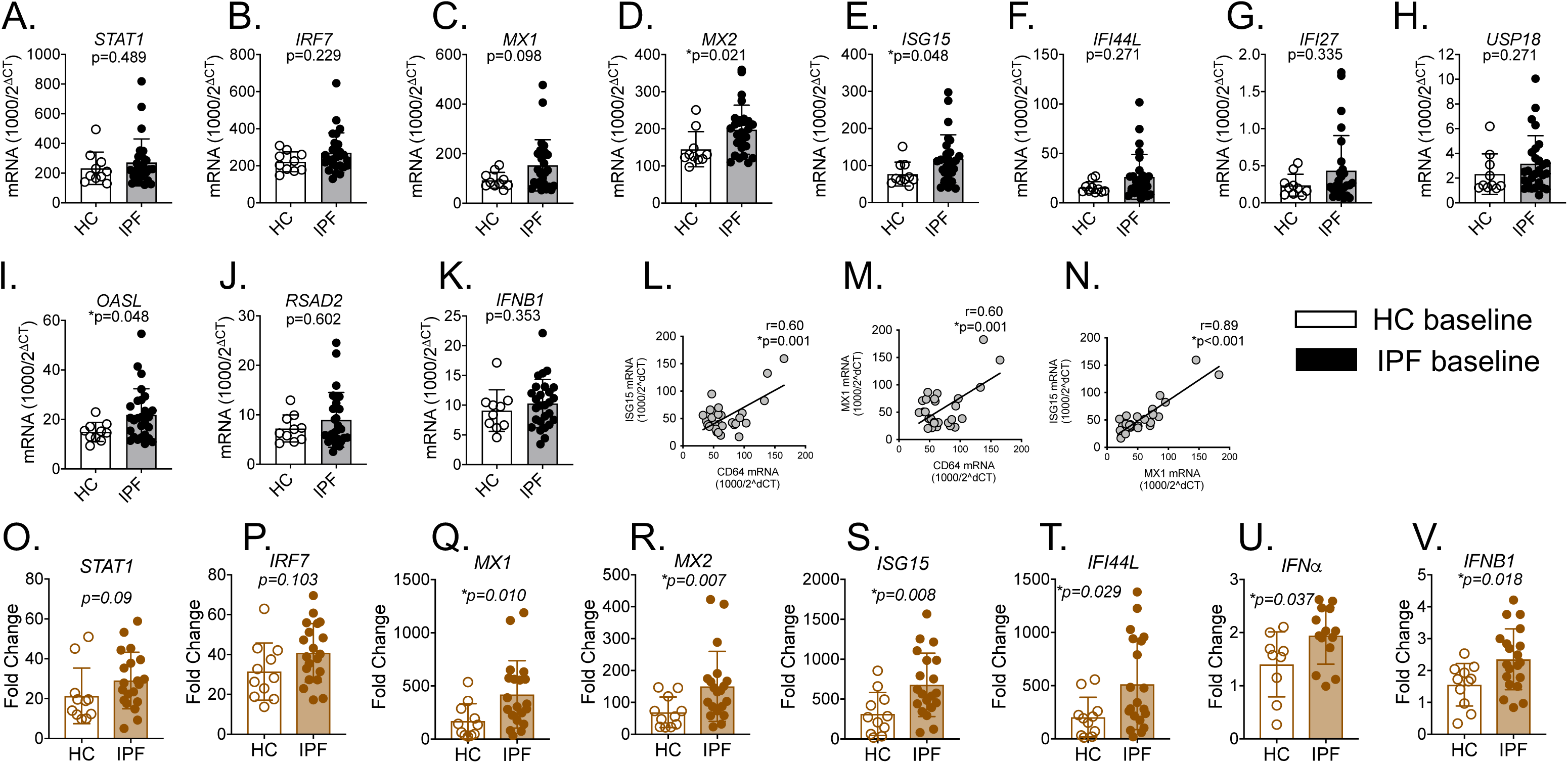
Increased interferon stimulated genes (ISG) expression and heightened response to type I IFN stimulation in monocytes from IPF. (A)-(L) ISGs, IFNB1 and IFNAR expression in freshly isolated IPF monocytes vs HC, using qPCR. Results were expressed as relative gene expression - 1,000/2^(CT of target gene−CT of housekeeping genes) (M-O) Correlation of CD64 mRNA expression levels in IPF monocytes with ISG15 and MX1 levels. (P-W) Increased levels of ISG mRNA expression in IPF monocytes after culture in recombinant type 1 IFN for 18 hours. Fold change refers to change over unstimulated values for that patient’s monocytes.

To explore whether CD64 expression were linked to type 1 IFN signalling, contemporaneous basal expression of CD64 *(FCGR1B)* and representative ISGs (*MX1* and *ISG15*) were also examined. In patients where CD64, MX1 and ISG15 gene expression in monocytes was measured concurrently, we observed a strong positive correlation between all three genes, supporting the link between CD64 expression and type 1 IFN signalling (Figure 4L-M). In this experiment, as an internal control, MX1 expression was tested against ISG15, and as expected, expression of these ISGs correlated positively and very closely with each other attesting to the rigour of the data (Figure 4N).

Next, we tested if monocytes showed an enhanced response to type 1 IFN stimulation. Freshly isolated monocyte from IPF patients (n=20) and HCs (n=10) (Supplemental Table 8B) were stimulated with 100U/mL recombinant human IFN-β for 18 hours and expression of key ISGs, *IFNA* and *IFNB1* were examined.

Upon stimulation with type I IFN, both IPF and HC, as expected, monocytes showed increased expression of ISGs when compared to unstimulated monocytes (Figure 4O-V). The increase was significantly greater in IPF monocytes for *MX1*, *MX2*; *ISG15* and *IFI44L* (p=0.010, p=0.007, p= 0.008 and p= 0.029 respectively) (Figure 4N-S). The induction of both *IFNA* (p=0.04) and *IFNB* (p=0.02) was also significantly greater in IPF monocytes (Figure 4U-V). These results are in keeping with a primed type I IFN signalling pathway, along the JAK-STAT signalling arm of the pathway (Supplemental Figure 3).

As with basal level changes, the expression of ISGs after IFN-β stimulation was also strongly correlated with each other. The strongest links were between stimulated *MX2* and *IFI44L; MX1* and *MX2*, and *MX1* and *IFI44L* (all p<0.001) (Supplemental Figure 4U), providing internal control for the data.

These findings suggest that IPF monocytes are primed to respond to type 1 IFN stimulation at homeostasis, and accordingly, respond in a heightened manner when cultured with recombinant IFN-β. Importantly, CD64 expression appear to be linked to basal type 1 IFN priming, supporting the suggestion that CD64 is a marker of the type 1 IFN signature found in IPF monocytes.

### Monocytes from IPF patients showed impaired differentiation to macrophages

In the final study, we evaluated the potential impact of our observed monocyte abnormalities in IPF patients on monocyte-derived macrophages (MDMs). Freshly isolated monocytes from IPF and HCs were differentiated to MDMs over 7 days using 10% autologous serum.

As CSF-1 has been shown to influence the differentiation of monocytes towards an M2 macrophage phenotype (39–41), we first examined whether IPF MDMs exhibited M2 features. We found no M2 phenotypic bias on MDMs after 7 days of culture – there was no difference in CD163 or CD200R expression on MDMs derived from IPF patients (n=17) compared to HCs (n=19) (Figure 5A-B). We also analysed the expression of CD64 on MDMs but found that both IPF and HC MDMs showed reduced levels of CD64 expression as they differentiate to MDMs (Supplemental Figure 5 C-D). Thus, the increased CD64 expression in IPF monocytes (compared to HC) appear a feature of monocytes rather than monocyte-derived macrophages.

**Fig 5.**
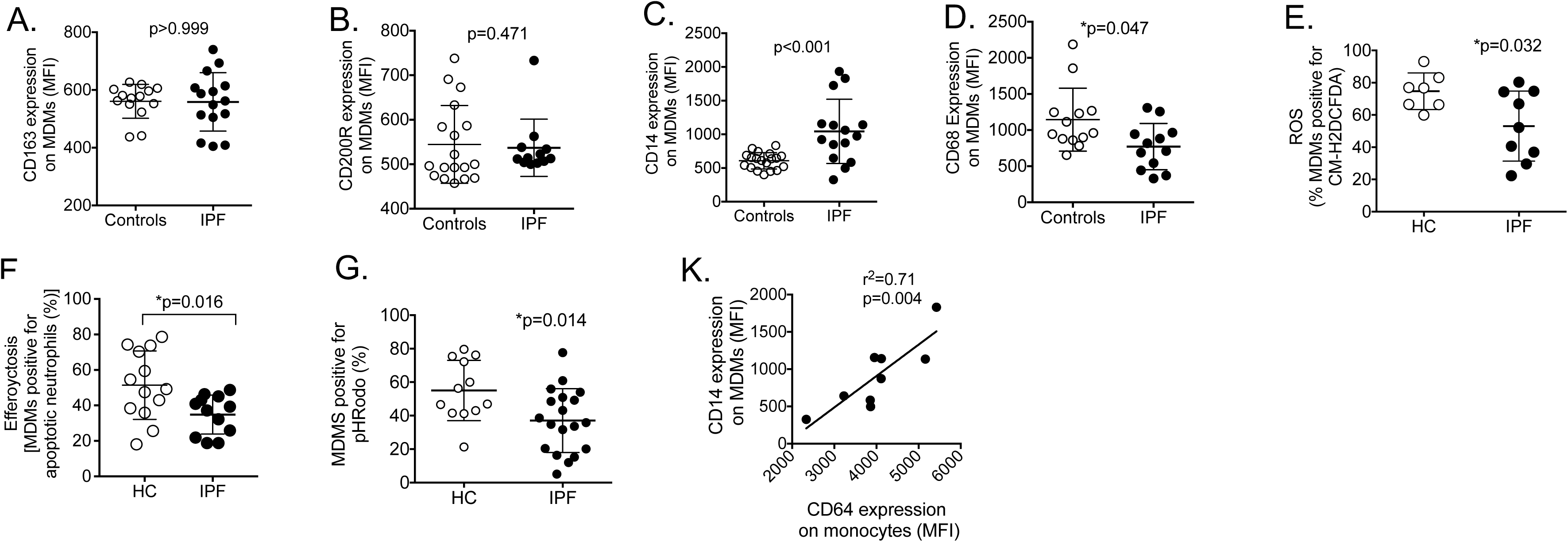
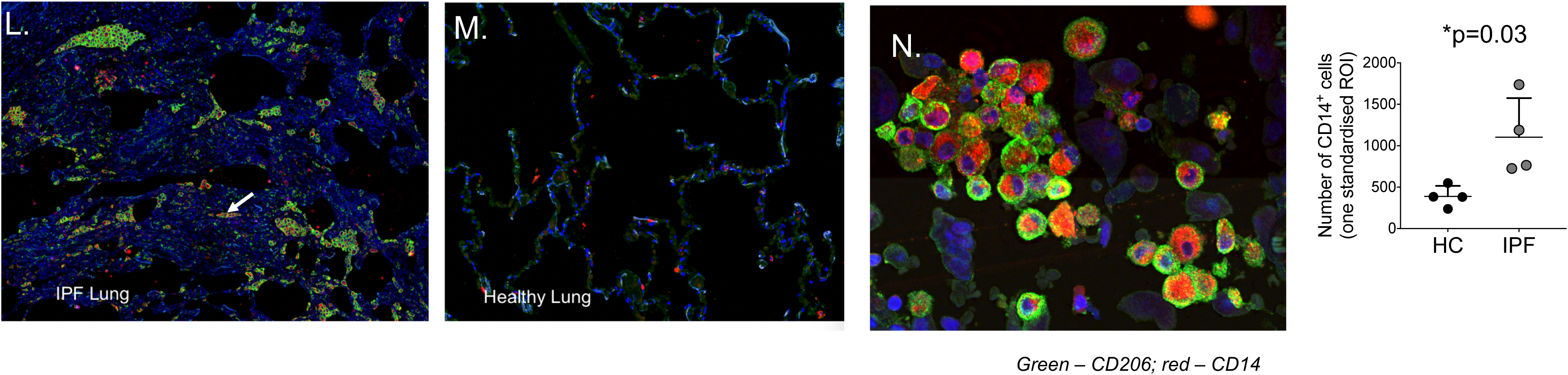
A-K. IPF monocyte-derived macrophages show impaired differentiation to macrophages. (A)-(B) flow cytometry expression of representative M2 markers(CD163 and CD200R) on monocyte-derived macrophages (MDM) after 7 days culture with 1% autologous serum. (C) –(D) Retention of CD14 and lack of expression of CD68 on IPF MDMs by flow cytometry on day 7 indicating poorer progression from monocytes to MDMs compared to monocytes derived from healthy controls (HC). (E) ROS production by MDMs after induction of oxidative stress using 0.03% H_2_O_2_ (F) Efferocytosis of aged neutrophils by MDMs. Neutrophils were labelled with a cell tracer, aged for 18-20 hours’ post venesection and co-cultured with MDMs at a ratio of 1:3 (neutrophils: MDMs) for 2 hours then fixed. Cell populations positive for the tracer dye were indicative of neutrophil efferocytosis by MDMs (G) Phagoyctosis capacity of MDMs using *pHrodo® Green E. coli* bioparticle assay. (K) Correlation between CD64 levels on monocytes and the ability to differentiate (the latter represented by CD14 expression levels on day 7 MDMs) (L)-(M) Representative immunofluorescence staining on lung sample from one IPF patient and one control. Macrophages are marked by CD206 expression (green), and found in groups lining the alveolar space of IPF but not non-IPF control lungs. (N) Higher magnification shows a typical area of macrophages which are co-expressing CD14 together with those which are CD14^-^ (O) Number of CD14 expressing cells in standardised area of region of interest (ROI) in lungs of IPF and controls (n=4 each).

However, on closer inspection, we observed a clear but unexpected impairment in capacity of IPF monocytes to differentiate into mature MDMs compared to healthy controls. After 7 days of culture, IPF MDMs retained expression of the monocyte marker, CD14 (Figure 5C), and expressed lower levels of CD68, a marker of fully differentiated macrophages (Figure 5D) compared to control MDMs from age-matched healthy controls (Supplemental Figure 5A-B). To further support these data, we analysed the ability of monocytes to differentiate into macrophages by examining three prototypical macrophage functions – (i) endogenous production of reactive oxygen species (ROS), (ii) efferocytosis and (iii) phagocytosis. In keeping with a less mature or differentiated macrophage phenotype, IPF MDMs showed significantly lower rates of the archetypal macrophage functions-cellular oxidant levels (Figure 5E), efferocytosis (Fig 5F) and phagocytosis (Figure 5G) (Supplemental Table 9 for demographics of recruited patients).

These results show that monocytes from IPF patients differentiated to macrophages less readily and therefore remained monocytes for longer. This is likely to have several implications in the repair, fibrotic and immune defence capacity in IPF (16). However, here, we were particularly interested in exploring if it might be linked to enhanced type 1 IFN signalling in IPF monocytes. To test this, we used monocytic CD64 expression as a reflection of type 1 IFN signalling (as suggested in Figure 4L-M), and MDM CD14 expression as marker of macrophage differentiation [Supplemental Figure 5A-B] and questioned if monocytes expressing high CD64 subsequently showed lesser ability to differentiate into macrophages (MDM with higher CD14 levels indicate more monocytic feature). We found this to be the case, CD64 expression on the initial monocytes strongly and positively correlated with the retention of CD14 on MDMs differentiated from these monocytes (Figure 5K).

To explore whether there was evidence to support persistence of monocytes in tissue, we examined the number of monocytes and macrophages in IPF and healthy controls. Biopsy samples from IPF lungs (n=4) were compared to non-fibrotic lung tissue taken from areas distant to resected lung margins in localised lung cancer with no lymph node spread or metastasis (Table S10 for demographic details). Monocytes and macrophages were identified using immunofluorescence staining as follows: monocytes were CD14^+^CD206^neg^; mature macrophages were CD14^neg^CD206^+^ and differentiating monocyte/macrophages were CD14^+^CD206^+^.

IPF lung samples showed a clearly expanded interstitium with large numbers of CD14^+^ and CD206^+^ monocyte/macrophages in the alveolar space (Figure 5L). This monocyte/macrophage population was not seen in non-fibrotic lungs (Figure 5M). Cells in the IPF samples appeared clustered together (Figure 5L), and on high magnification, were revealed to be a combination of monocytes (CD14^+^CD206^-^), maturing macrophages (CD14^+^CD206^+^) and mature macrophages (CD14^-^CD206^+^) cells (Figure 5N). To quantify CD14 expressing cells, an application was created using the Visiopharm Integrator System® platform where a standardised area of each lung slide was analysed for CD14-expressing cells (Supplemental Figure 6A-D) using pre-defined optimised machine-based algorithm (see Methods). This method did not differentiate between monocytes (CD14^+^CD206^-^ cells) and differentiating macrophages (CD14^+^CD206^+^ cells). However, it was clear that CD14 expressing cells (representing both monocytes and differentiating monocyte/macrophages) were significantly higher in IPF compared to control lungs [1104 (+/- 471) vs 433 (147) cells in 0.25cm^2^; p=0.03] (Figure 5O).

Taken together, these studies suggest that IPF monocytes have decreased capacity to differentiate to monocyte-derived macrophages and this is supported by presence of higher numbers of CD14^+^ cells in the lungs.

## DISCUSSION

Our results show for the first time, that IPF monocytes are endowed with a type 1 IFN gene expression signature and a primed type I IFN pathway, denoted by a heightened response to type 1 IFN stimulation *ex vivo*. These monocytes also displayed impaired capacity to differentiate to macrophages *ex vivo*, indicating increased longevity in survival as monocytes, in the lungs. These abnormalities appear linked to each other – monocytic ISG expression at baseline correlated significantly with expression levels of CD64, which in turn is inversely linked to the monocytes’ ability to differentiate to macrophages (Figure 4L-M, and 5K respectively). The frequency of circulating monocytes correlated with the severity of lung fibrosis, while the serum environment (with a markedly increased CSF-1 levels) supports the maturation of pro-monocytes to monocytes in the bone marrow and the survival of monocytes (29).

Our findings are important from several perspectives. Firstly, we provide a biological basis for the involvement of monocytes in the pathogenesis of IPF, and therefore stronger justification to explore these cells as potential therapeutic target beyond increased numbers of monocytes shown by both ours and Scott et al’s study (42). Our work also moves the mechanistic focus away from the pathological site of disease (lungs), to upstream pathways (in the blood) that may be contributing to fibrotic processes, potentially implicating the bone marrow, where monocytes are generated.

Enhanced responsiveness to type 1 IFN in monocytes is a highly significant finding for IPF patients. Type 1 IFN signalling is activated by two consecutive pathways. Sensing of self or viral nucleic acid and bacterial pattern-recognition molecular patterns (PAMPS) by TLR4 on the cell surface, or TLR3,7,8,9 in the endosome and other cytosolic sensors (Supplemental Figure 3) results in transcription of type 1 IFNs and some ISGs (31, 32). The type 1 IFN (endogenously produced by the cell via this pathway or from other cells) then ligates the IFN receptor (IFNAR) and results in hundreds of ISGs which directly interfere with pathogen replication and also activates other immune pathways to combat infection (31, 32). This highly potent immune defence pathway unleashes a large number of cytokines, chemokines (43–45), and also enhances natural killer cell function and high affinity T and B cell responses (46). In non-infectious settings, Type 1 IFN signalling can be persistently activated [e.g. due to the recognition of self-nucleic acid in SLE and sensing of abnormal nucleic acid species in Aicardia-Goutieres syndrome (AGS) resulting in severe inflammation and clinicopathologic manifestations (35)]. In health, type 1 IFN signalling is tightly regulated, and its activation is accompanied by restraining pathways to reduce the impact of inflammation and immune activation (46, 47). Our findings show that IPF monocytes are ‘held’ in a heightened state of type 1 IFN signalling readiness, with the ability to respond at greater magnitude to type 1 IFN stimulation, which may in turn result in over-exuberant downstream innate and adaptive immune responses. This is an important finding in IPF as it suggests that monocytes have greater capacity to cause injury during an infection or when stimulated by damage-associated molecular patterns (DAMPS) (76). The former trigger is particularly relevant in AEIPF (Figure 1H), a devastating event associated with diffuse alveolar injury, accelerated fibrosis and high mortality. The cause of AEIPF is unknown but often associated with an infective trigger (48). Trafficking of large numbers of monocytes to the lungs can precipitate injury due to release of inflammatory mediators like IL-6, TNF-α and IL-1β (49). In IPF, injury could be further enhanced by ‘super-charged’ monocytes primed to respond with an augmented type 1 IFN response. In fact, the diffuse alveolar damage pattern found in lungs during AEIPF (on both histology and CT imaging assessment), is akin to the widespread alveolitis in high-pathogenicity influenza infection where type 1 IFN and monocytes are central causal factors. In a lung microenvironment set for aberrant repair, as in IPF, such injury could lead to the accelerated fibrosis observed in AEIPF, and drive progression of fibrosis as seen in IPF. Delayed differentiation of monocytes to macrophages could mean persistence of pro-injury monocytes in the lung. Indeed, monocytes per se are recognised mediators of inflammation and injury. In other organs (e.g. neuronal damage models), the accumulation of monocytes is associated with extensive secondary axonal death and delayed repair (50). Similarly in the liver, monocyte presence is associated with pathological inflammation (51). We have also shown previously that very high levels of circulating monocytes are associated with severe respiratory impairment in influenza (52). Reduced rate of differentiation to macrophages can also have other negative impact on fibrosis. Macrophages are important effector cells in mediating repair and limiting inflammation after the initial stages of injury. They scavenge cellular debris, dying neutrophils and other apoptotic cells, and release IL-10 and other factors that regulate and control extracellular matrix deposition (6, 53). It is unclear why IPF monocytes showed poorer differentiation capacity but correlation with monocytic CD64 expression suggests that this may be related to enhanced type 1 IFN signalling. Intriguingly, a murine study that showed type 1 IFN activation in infiltrating monocytes delayed their differentiation to macrophages in the liver during hepatitis C infection, although the mechanism for this was unclear (54).

Our study did not elucidate the cause of type I IFN signalling priming. We did not observe high levels of IFN-α or β in the serum nor in the transcriptome of IPF monocytes. This could be due to well recognised difficulties in measuring type 1 IFNs at the RNA and protein level, and if so, it is possible that low level type 1 IFN production occurs in IPF patients. Against this possibility is the selection criteria for our patients, which rules out over infection and minimises the chance of subclinical infection. Clinical assessments were undertaken to ensure that there were no symptoms of infections nor evidence of recent clinical deterioration. However, IPF patients have been shown to have greater levels of latent viral particles, e.g. Herpes and Epstein Barr viruses in lung biopsies (55–57). Although these viruses tend to be kept in check by T cells, there is evidence that type I IFN signalling may also be involved (58), so this could potentially be a pathway to a primed type I IFN signalling in these patients. The other possibility is an epigenetic influence on the transcription of type I IFN signalling. The interferon regulatory factors (IRFs) are under epigenetic control (59) and can be modulated epigenetically to increase type I IFN signalling. These possibilities will be interesting to explore in future studies.

Increased CD64 expression on monocytes was a key feature of IPF monocytes. This was the only clear phenotypic abnormality that we observed, and its level of expression correlated with serum CSF-1 levels, ISGs (MX1 and ISG15), and the persistence of monocytes in the *in vitro* studies; linking these findings together. These links and it being a surface protein make increased monocytic CD64 a potentially useful marker for the abnormalities observed in IPF monocytes. CD64 is a high-affinity receptor (FcγRI), which binds the Fc portion of IgG, and is constitutively expressed on monocytes and macrophages. It has also been shown to be an ISG (60, 61). CD64 on monocytes has been observed to correlate with IFN-α levels in the serum of SLE patients and enhanced type 1 IFN signalling (37). Selective apoptosis of CD64^hi^ macrophages in skin inflammation model was associated with resolution of skin inflammation (62). Of relevance to our findings, using type 1 IFN knockout mice, Fleetwood and colleagues showed that type 1 IFN signalling significantly enhanced CSF-1 production in murine macrophages (41). Thus the correlation between CD64 and CSF-1 levels (Supplemental Figure 2K) could reflect the correlation between type 1 IFN signalling and CSF-1 production. Our studies do not confirm this however and more work is required to test this hypothesis.

Collectively, our data implicate a ‘super-charged’ monocytes as a potential driver in IPF, possibly by enhancing alveolar epithelial injury through the persistence of monocytes with a potentially injurious, primed type I IFN signalling in the lungs. Higher CD64 expression could be secondary to this enhanced type 1 IFN signalling, and the latter could also be the cause for the delayed macrophage differentiation. Whilst the data do not establish a causative relationship between the observations, these novel findings provide an impetus to move the mechanistic focus to immune pathways outside the lungs. Inhibiting or modulating CD64 expression could be a therapeutic possiblity in IPF, given the selective expression of CD64 on myeloid cells, and its high affinity for IgG, which makes antibody-mediated inhibition an attractive strategy The findings also emphasise the importance of infection, in driving progression of, and precipitating widespread damage and acceleration of fibrosis in IPF.

In summary, our study identifies and defines the key monocytic abnormalities in IPF, describing type 1 IFN primed monocytes as possible mediators that exacerbate injury which worsens an already aberrant repair process. We have identified increased levels of CD64 as a potential new theragnostic target for IPF. These findings provide fresh mechanistic insight into a devastating disease with limited therapeutic options.

## Methods

**Patient and control recruitment, and imaging methods in Supplemental Methods.**

### Isolation of PBMCs and monocytes

Blood samples were collected in lithium heparin (Greiner Bio-one) and processed within 4 hours. Peripheral blood mononuclear cells (PBMCs) were extracted by Lymphoprep^TM^ (Axis-Shield) density gradient separation. Monocytes were isolated from PBMCs by positive selection using anti CD14 microbeads (Miltenyi Biotec) according to the manufacturer’s instructions. Post separation CD14^+^ monocyte purity was assessed by flow cytometry (CD3, CD19, CD15, CD16 and CD14). Only samples with purity of greater than 98% were used.

### Serum preparation and soluble mediator analyses

At the time of sampling, all participants had blood taken for serum using SST Vacutainer® serum separator tubes (BD), and serum isolated as per manufacturer’s instructions and stored at −20°C for batch testing. CCL2 (MCP-1), IL-6, CXCL-10 (IP-10), TNF-α, IFN-γ, CSF-2 (GM-CSF), CCL-20 (MIP3A), CXCL-8 (IL-8), IL-10, IL-13, IL-1*b*, and IFN-*b* levels were measured in serum by human magnetic bead Luminex multi-analyte assay following the manufacturer’s protocol (Bio-techne custom analyte mix). Results were obtained with a Bio-Plex 200 System (Bio-Rad). CSF-1 was measured by standardized sandwich ELISA (R&D Systems).

### Flow Cytometry

All antibodies were purchased from Biolegend, except Annexin V (eBiosciences). Cultured cells were incubated with Zombie Aqua fixable viability stain (Biolegend) to enable exclusion of dead cells from subsequent analysis. Cells were incubated with antibodies against surface antigens for 20 minutes at 4°C and fixed with Stabilising Fixative (BD) prior to acquisition (52). For detection of CD68, cells were permeabilized with saponin (Sigma-Aldrich) and incubated with anti CD68 for 30 minutes at room temperature. Cells were acquired using a LSRFortessa™(BD) and data was analyzed using Flowjo v10 software (Tree star, Inc) and FACSDiva™(BD).

### IFN-β stimulation of cultured monocytes

CD14 monocytes plated at 200,000 cells per well in 96well flat bottom plates in 200μl complete RPMI-1640 (R10) media supplemented with 2mM L-glutamine, 100IU/mL penicillin/streptomycin and 10% heat-inactivated fetal calf serum (Sigma-Aldrich). Monocytes were stimulated for 18hrs with 100U/mL recombinant human IFN-β (R&D systems) and processed for RNA extraction.

### Real-time PCR

For all qPCR phenotypic studies (Supplemental Table 4 and Figure 4), CD14 positively-selected monocytes were lysed in RLT buffer (QIAGEN) supplemented with beta-mercaptoethanol (Sigma-Aldrich) before RNA extraction using the RNeasy Mini Kit (QIAGEN) and frozen at −80°C. Batch RNA extraction was performed using the RNAeasy Mini Kit (Qiagen) according to manufacturer’s protocol. RNA integrity was assessed using nanodrop and Agilent technology. For data in Supplemental Table 4, cDNA was synthesized using High Capacity cDNA RT Kit (Applied Biosystems). qPCR was performed using the 2X Fast SYBR® Green Master Mix (Applied Biosystems) in a 7500 Fast Real-Time PCR system (Applied Biosystems). Primer sets are shown in table S11 in Supplemental Methods.

For qPCR type 1 IFN signalling studies (Figure 4), real-time PCRs for genes were performed using TaqMan Fast Advanced Master Mix (Applied Biosystems) with TaqMan primer/probe sets for human genes [*STAT1, IRF7, MX1, MX2, ISG15, IFI44L, IFI27, USP18, OASL, RASD2, IFNB1, IFNA1*, and *FCGR1A* (CD64)]. Real-time PCRs were performed on a QuantStudio 7 Flex Real-Time PCR System and threshold cycle (CT) values were determined from duplicate reactions using QuantStudio software (Thermo Fisher Scientific).

Results were expressed as relative gene expression = 1,000/2^(CT of target gene−CT of housekeeping genes) when there were no start or control group (63) ie when measuring basal gene expression (Figure 4A-K) and fold change relative to unstimulated group (Figure 4N-U).

### Bulk RNA sequencing

3 IPF patients (all male; aged 57,76 and 78y) and three healthy controls (HC) (all male, non-smoker, no medications; aged 65,68 and 71y) were recruited from the Oxford Interstitial Lung Disease Service. Circulating monocytes were isolated by CD14^+^ Macsbead positive selection and RNA extracted using RNeasy Mini Kit (Qiagen) as described above. RNA integrity number (RIN) exceeded 9 for all samples measured by 2100 Bioanalyser (Agilent). RNASeq libraries where prepared using Smartseq2 as described by Picelli et al (64). Briefly cells were sorted into lysis buffer and converted into cDNA. Resulting libraries were converted to Illumina compatible libraries using Illumina Nextera XP kit, as manufacturers instructions, and sequenced using NextSeq 500(Illumina) (single end, 75 bp unpaired sequencing). Sequencing depth was 30 million per sample.

FASTQ files were generated and quality of raw sequencing reads was initially assessed using fastQC (65). Poor quality bases (<20) and technical sequences were trimmed using Cutadapt software and reads were subsequently aligned using STAR aligner (66) against the human genome hg38 assembly. Non-uniquely mapped reads were discarded and gene expression levels were quantified as read counts using the featureCounts function (67) from the Subread package (68) with default parameters. There were 27.4-38.3M final uniquely mappable reads per sample. The read counts were normalized using library size factors to account for differences in sequencing depth and/or RNA composition, calculated using the median ratio method, as described in Anders and Hubert (69). Differential expression performed using the DESeq2 R package (70). R package clusterProfiler (71) was used to perform enrichment studies on differentially expressed genes and GSEA (30) on all genes between the two groups, using gene sets from GO (72–74) and Reactome (75) pathways. Significance threshold was set at 5% FDR in all cases.

### Generation of monocyte-derived macrophages (MDMs)

Monocytes were cultured in X-VIVO^TM^ (Lonza) supplemented with 10% autologous serum. Cells were plated onto low-adherence plates (Corning) and cultured for 7 days. 50ng/ml human M-CSF (Miltenyi Biotec) was added on day 0 to promote monocyte survival. On day 7, plates were placed on ice for 20 minutes to detach monocytes which were then harvested by gentle pipetting, washed and counted.

### Phagocytosis assay

pHrodo® Green E. coli bioparticle assay (Molecular Probes) was used according to manufacturer’s instructions. Upon phagocytosis and transfer into the lysosome, the pH sensitive bioparticle emits a fluorescent signal. MDMs were incubated with pHrodo® Green E. coli bioparticles for 30min at 37°C, washed stained with viability dye Zombie Aqua (Biolegend), placed on ice before immediate flow cytometry analysis.

### ROS assay

CM-H2DCFDA (Molecular Probes) was used according to manufacturer’s instructions as indicator for reactive oxygen species (ROS) in cells. MDMs were cultured for 7 days as described above and incubated with 5 µM CM-H2DCFDA at 37°C for 30min. Oxidative stress was induced using 0.03% H_2_O_2_ for 1 hour then cells were washed stained with viability dye Zombie Aqua (Biolegend) and placed on ice before immediate flow cytometry analysis.

### Neutrophil efferocytosis assay

Neutrophils were isolated from whole blood using the MACSxpress isolation kit (Miltenyi Biotec), according to the manufacturer’s instructions, and then stained with the cell tracer Far Red (Molecular Probes) and incubated overnight at 37°C. MDMs were stained with cell tracer Violet Proliferation Dye (Molecular Probes) and aged neutrophils (between 18-20 hours post venesection) were then added to MDMs at a ratio of 1:3. Cells were incubated at 37°C for 2 hours then fixed with Stabilising Fixative (BD). Co-cultured cells were then examined by flow cytometry and cells positive for both tracer dyes were indicative of neutrophil efferocytosis by MDMs. Cytochalasin D (Thermo Fisher Scientific) which inhibits the process of phagocytosis, was used as a negative control.

### Human lung Immunofluorescence

Paraffin-embedded human lung tissue sections [The Oxford Centre for Histopathology Research, OCHRe Oxford Radcliffe Biobank (ORB)] were deparaffinized and each section was pre-treated using heat mediated antigen epitope retrieval and stained with anti-CD14 (Abcam, AB183322) and Anti-MRC1/CD206 (Atlas Antibodies, AMab90746) overnight at 4°C. These were subsequently co-stained with Alexa Fluor 568 conjugated Goat Anti Rabbit IgG or Alexa Fluor 488 conjugated goat anti-mouse IgG2b secondary antibody and imaged using a Nikon Ti2 microscope (Nikon Instruments) attached to a Dragonfly 200 spinning disk confocal microscope (Oxford Instruments). Full protocol in Supplemental Methods.

Visiopharm Integrator System software (version is 2018.4.3.4480 Visiopharm) was used for quantification of CD14-expressing cells in human lung samples. Image analysis protocols are implemented as Analysis Protocol Packages (APP) in VIS. An APP was designed to quantify slides briefly, red colour band (RGB-R) is used to detect positively stained cells on red fluorescent slides with polynomial blob filter to enhance the edges of the cells. The RGB-R is input to a threshold classifier. As post-processing steps; a method for cell separation which is based on shape and size is used, cell areas that are too small are removed and finally applying unbiased counting frames to avoid the cells that are intersecting with neighbouring tile boundaries counted twice (or more).

### Statistical analysis

GraphPad Prism version 7.00 (GraphPad) or R version 3.4.3 were used for all statistical analyses. D’Agostino–Pearson omnibus normality test was used to determine the distribution of data where relevant. Unless described specifically in the texts, for comparison of data sets that showed normal distribution, the Student t-test was used. Mann-Whitney test was applied for data that was not normally distributed. To compare multiple data sets that were normally distributed, one-way ANOVA with Tukey’s correction for multiple comparisons was used. Multiple data sets that were not normally distributed were analysed using Kruskal-Wallis test with Dunn’s correction for multiple comparisons. Paired data sets were analysed using the paired t-test or Wilcoxon test when data did not show a normal distribution. For significance of correlation tests, data was Box-Cox transformed (to transform non-normal variables into a normal form) after which Pearson correlation testing was performed. A p-value of less than 0.05 was considered significant.

### Study approval

Patients were recruited from the Oxford Interstitial Lung Disease Clinical Service and the study had ethical approval from the local and UK national ethics committee (14/SC/1060 from the Health Research Authority and South Central National Research Ethics Service). Written informed consent was received from participants prior to inclusion in the study.

## Supporting information

All Supplemental Data

## ACKNOWLEDGEMENTS

The research was funded by the National Institute for Health Research (NIHR) Oxford Biomedical Research Centre (BRC) and Oxford-UCB Alliance research grant. LPH was part funded by the MRC(UK), EF and LD were funded by the Oxford NIHR Biomedical Research Centre (BRC). Training, support and use of the Visiopharm software platform were supported by the Oxford NIHR Biomedical Research Centre (Molecular Diagnostics Theme/Experimental Pathology sub-theme) Cancer Research UK (CR-UK) grant number C5255/A18085, through the Cancer Research UK Oxford Centre and the Pathological Society of Great Britain and Ireland.

We thank Dr Jennifer MacLellan for help with patient recruitment for the type I IFN studies. We thank the late Prof Vincenzo Cerundolo for scientific discussions, and the paper is in memory of him and his enthusiasm for the project.

## Authors contribution

EF performed all the experiments on monocyte and MDM phenotyping, MDM functional studies and recruited nearly all the patients. LD performed the type I IFN studies and organised recruitment of patients, KB performed the qPCR studies on monocytes, and the bulk RNA studies and contributed intellectually to the analysis, CV performed all the immunofluorescent studies, organised patient samples for type I IFN studies, AA and ER performed bioinformatic analysis on the bulk RNA sequencing data, contributed intellectually to the analysis and contributed to statistical analyses of all the data, VI provided statistical analysis and overview of the data and statistical handling of the serum profiling output, ST provided statistical overview of the study, NA performed the chemistry studies for the bulk RNA sequencing, VSN and RB were specialist thoracic radiologists who helped design and analysed the high resolution CT scans for the patients and quantified the extent of fibrosis, RH assessed patient suitability, and contributed to recruitment of patients, CC contributed to acquisition and analysis of lung samples, CH performed the Luminex studies and contributed to analysis, NKM performed the scoring of the CD14 cells in lung samples and with LPH, developed the application for quantification of the CD14 cells, RE and JR contributed intellectually, and to the design and methodology of the type I IFN studies, JD and YZ contributed to the intellectual discussion of the monocyte phenotyping studies, LPH conceived and led the analysis for all aspects of the study, supervised the experiments and drew together the results and manuscript. All authors contributed to the analysis of their respective part of their studies and reviewed the entire manuscript and its conclusions. EF and LD contributed substantially to the writing of the manuscript.

