## Supplementary material for "Type-1 IFN primed monocytes in pathogenesis of idiopathic pulmonary fibrosis": All Supplemental Data

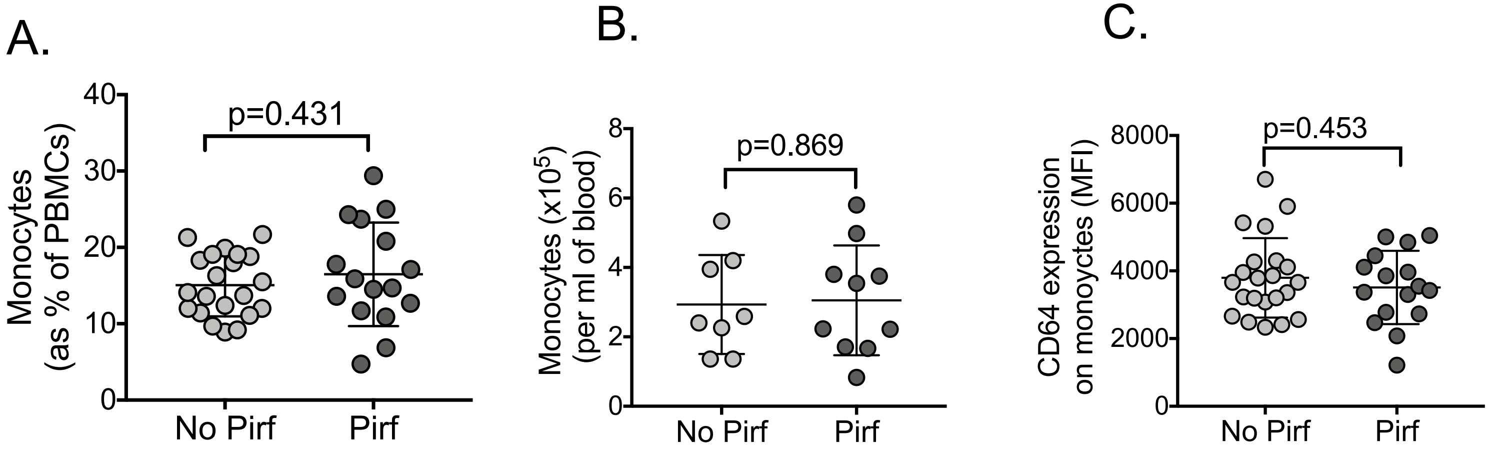


**Supplemental Fig. 1A-C** Monocytes levels and CD64 and CD163 expression levels on monocytes by flow cytometry in patients on Pirfenidone compared to without. MFI – mean fluorescence index. **(D)** qPCR of relevant genes in monocytes from IPF vs age-matched healthy controls, shown in graph form. Expression values normalized to three house keeping genes. Note both IL-6 and IL-10 showed trend towards increased values in IPF but no clear M1 or M2 polarization, nor pro or anti-inflammatory genes or cytokines expression.

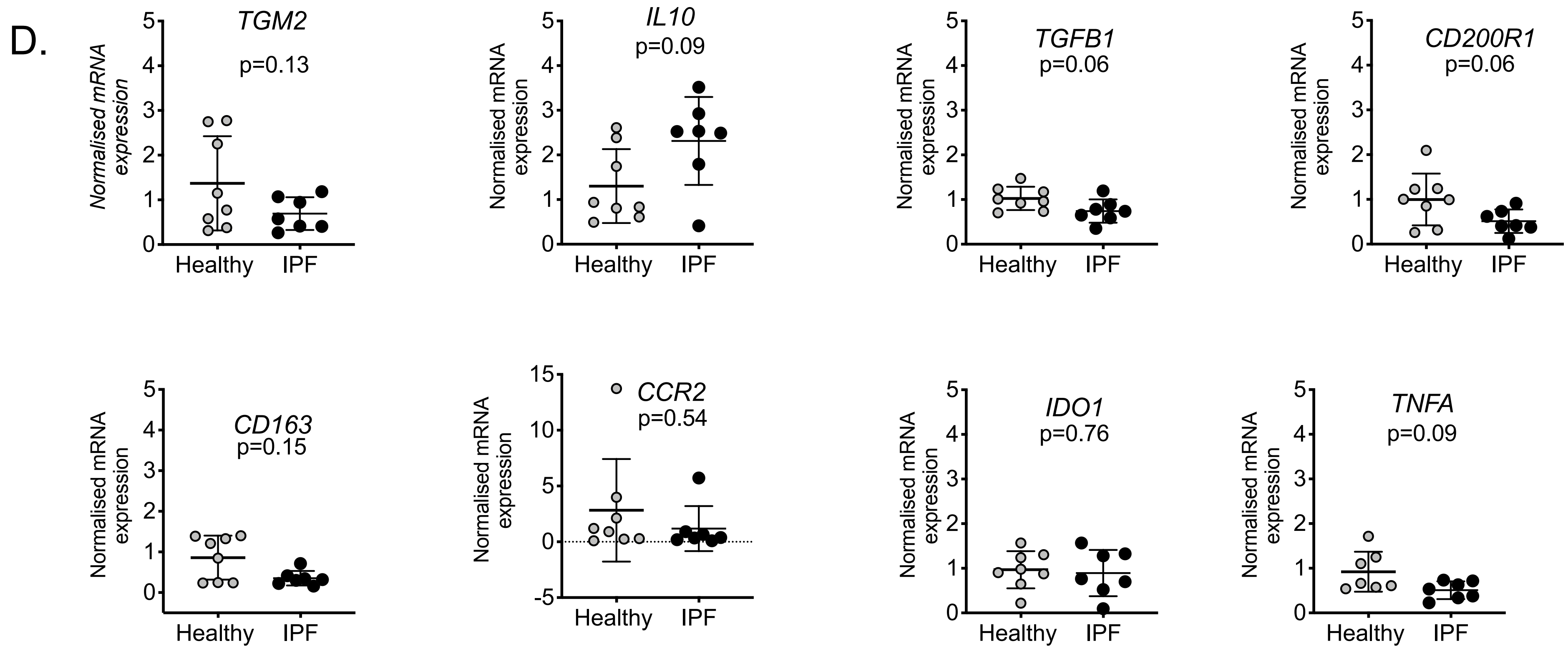


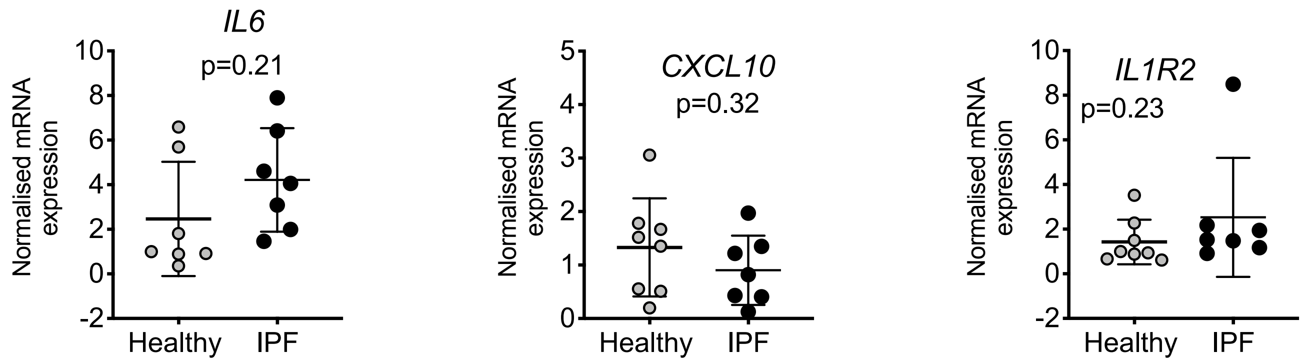


**Supplemental Fig 1D.** qPCR of relevant genes in monocytes from IPF vs age-matched healthy controls, shown in graph form. Expression values normalized to three house keeping genes. Note both IL-6 and IL-10 showed trend towards increased values in IPF but no clear M1 or M2 polarization, nor pro or anti-inflammatory genes or cytokines expression.

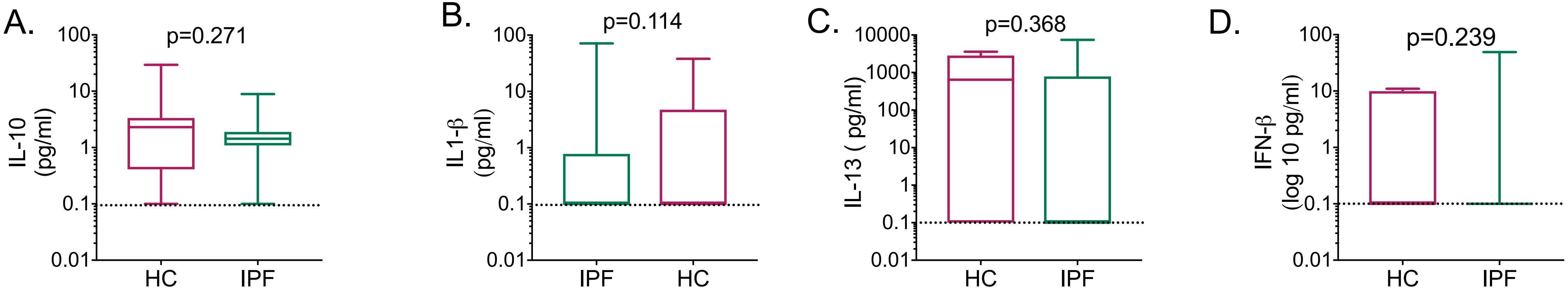


**Supplemental Figure 2A-D** Serum levels for IL-13, IL-10, IL-1β and IFN-β in IPF vs healthy controls (HC).

E.

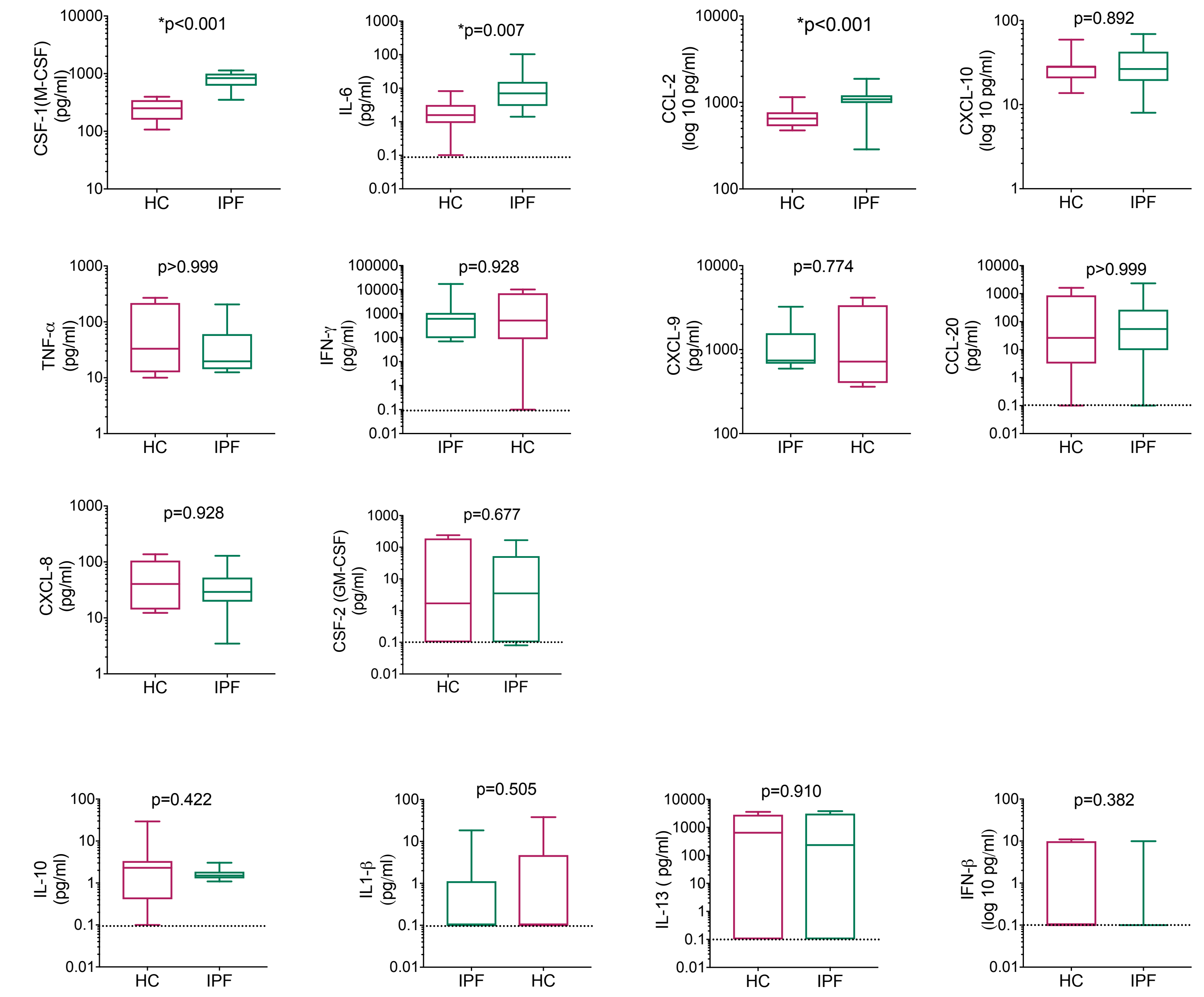


F.

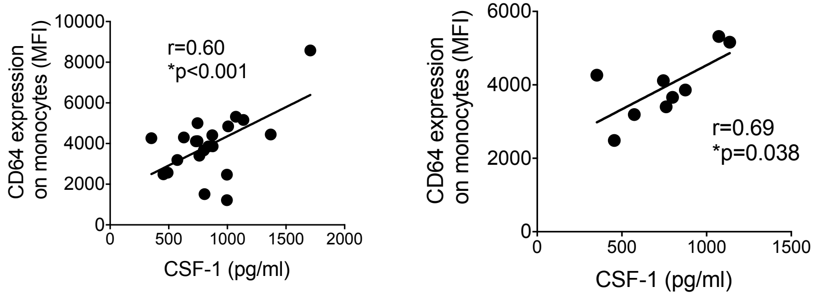


**Supplemental Figure 2 E-F. Serum profile of IPF patients –** mediators measured in serum as in Figure 2 but without patients on treatment (Pirfenidone or NIntedanib). Y axis is log 10. Zero values for serum mediators converted to 0.1 for visualisation purposes on a log 10 axis. Box plot is mean and S.D, whiskers reflect minimum and maximum values. P values calculated using Mann Whitney Rank Sum testing. (F) Correlation between serum CSF-1 levels and CD64 expression on monocytes (Pearson correlation); for all patients (left panel) and for those not on anti-fibrotics (right panel).

G.

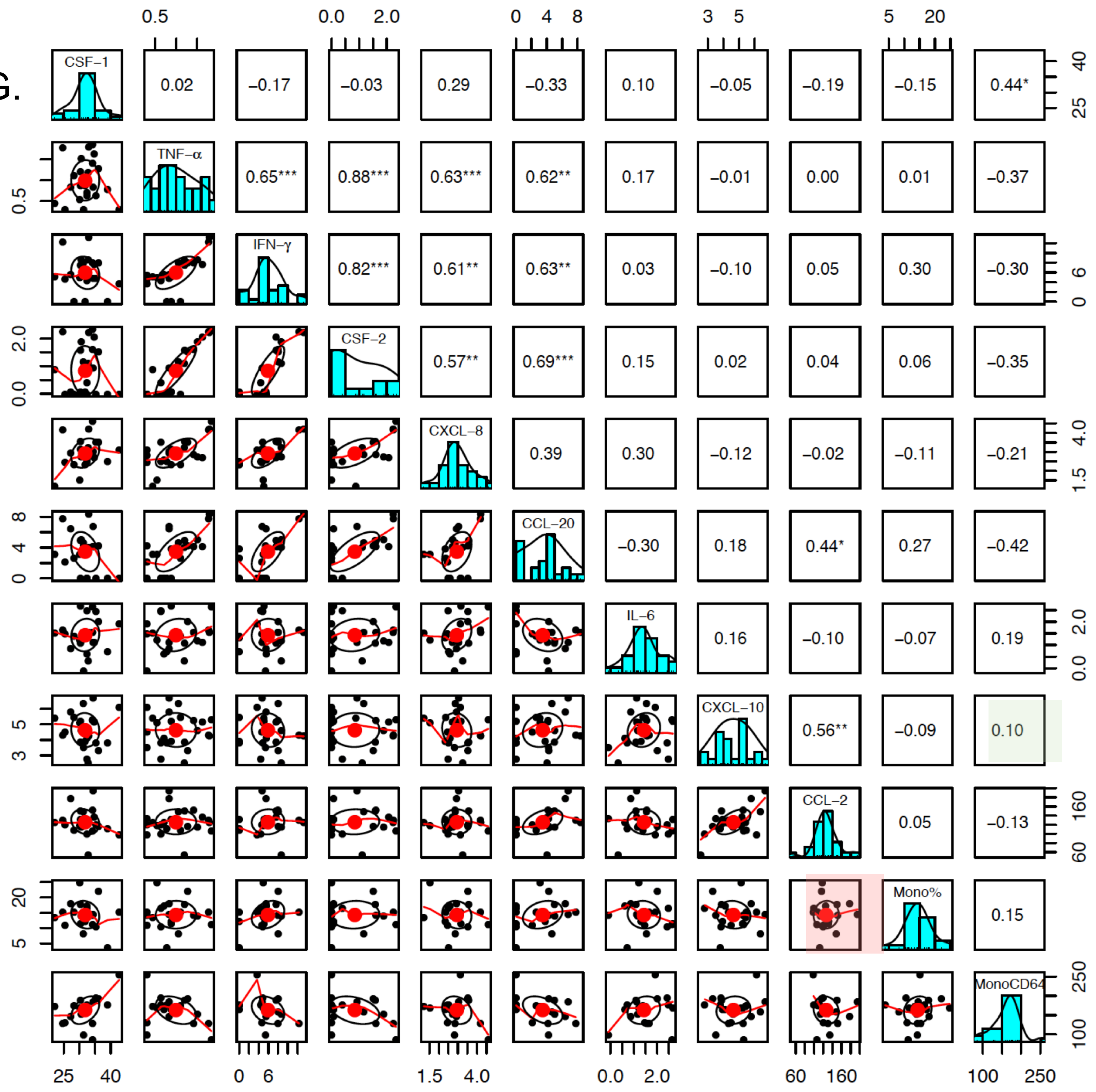


**Supplemental Figure 2G.** Pairs plot showing correlation between levels of all soluble mediators, monocyte levels and CD64 expression levels on monocytes. Values in boxes in right hand triangle refers to r value; graphs in boxes in left hand triangle displays scatter plots for corresponding correlation e.g. red box refers to correlation between CXCL10 and monocyte levels; r value is -0.09 (green box) and there was no significance. **p<0.001; *p<0.05. Bar histogram shows distribution of values for the named parameter. Only values from IPF patients are analyzed. (F) Amongst the soluble mediators, there was a strong positive correlation between mediators that might be secreted by the same cells eg CCL-2 and CXCL-10 from activated monocytes, and IFN-γ , TNF-α, CXCL-8, and CCL-20 which may be secreted lymphocytes. None of these showed different levels comparing IPF and healthy controls. Pearson correlation used for analysis after Box-Cox transformation $[(y^a-1)/a]$. With regards to non-normality of some variables, quantile normalizing the data did not change the conclusions.

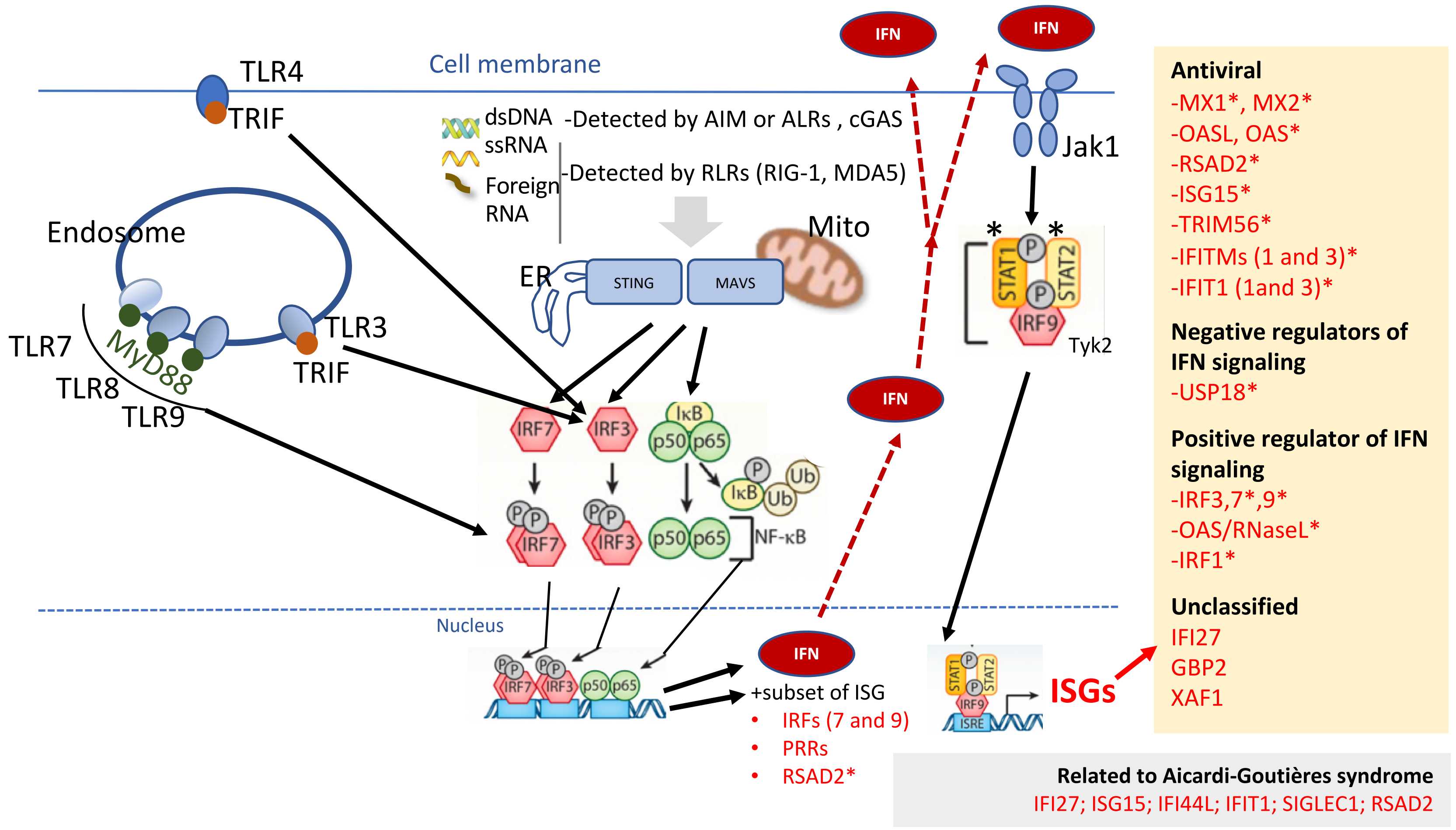


**Supplemental Figure 3. Abbreviated type I IFN signalling.** The endosomally expressed TLR-3, -7, -8, and -9, cell surface expressed TLR4, the RLRs [retinoic acid-inducible gene I (RIG)-I and MDA-5], and cGAMP synthase (cGAS) can couple pathogen (viral and bacterial PAMPs) and endogenous DNA (PAMPs and DAMPs) detection to type I IFN induction. TLR3 and TLR4 signal *via* TRIF, which occurs through inhibitor of kappa-B (IκB) kinases (IKKs), tumour necrosis factor (TNF) receptor-associated factor (TRAF) family associated NF-κB activator (TANK)-binding kinase-1 (TBK1), and IKK-ε (not shown in figure). This causes the activation of IRF3, which in turn induces the expression of type I IFNs. The activation of TLR3 can also induce the production of inflammatory mediators. TLR7, 8, and 9 use MyD88 for downstream signalling and can activate IRF and NF-κB pathways. RIG-I and MDA5 signal through the adaptor molecule mitochondrial antiviral signalling protein (MAVS). cGAS signals *via* the adaptor protein stimulator of interferon genes (STING). MAVS and STING further recruit signalling molecules (involving the IKK complex, TBK1, and several TRAF proteins) and lead to the activation of NF-κB and IRF3, resulting in gene expression of various antiviral cytokines including type I IFNs. Asterisks denote genes found in leading edge genes from GSEA analysis of monocyte RNA sequencing data (Fig. 3D-F).

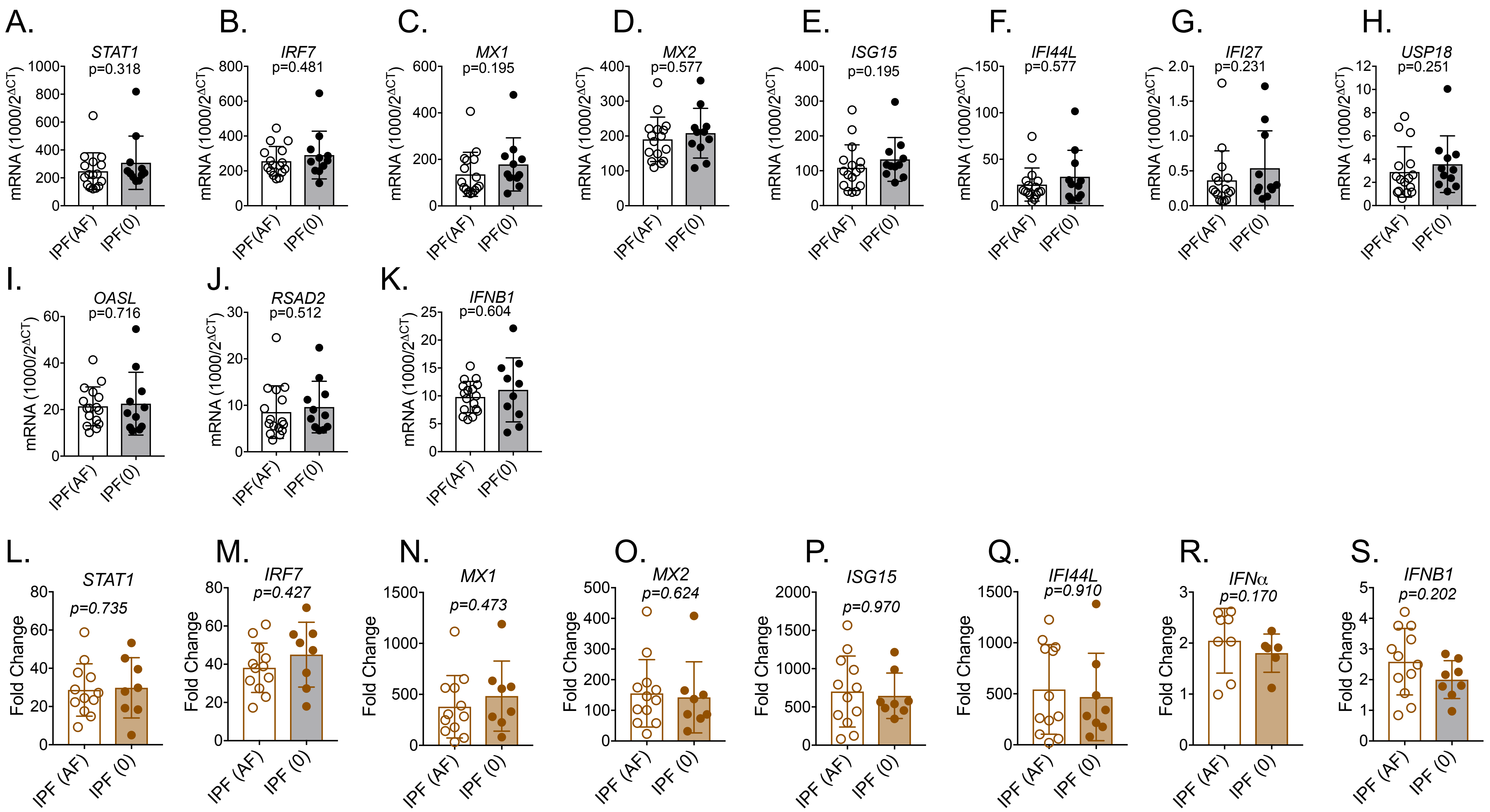


**Supplemental Figure 4A-S Interferon stimulated genes (ISG) expression at baseline** (A)-(K) and after culture in recombinant type 1 IFN for 18 hours ((L-S), comparing IPF patients who are on antifibrotics (Nintedanib or Pirfenidone) ('AF') and those not on antifibrotics '0'. Fold change refers to change over unstimulated values for that patient's monocytes.

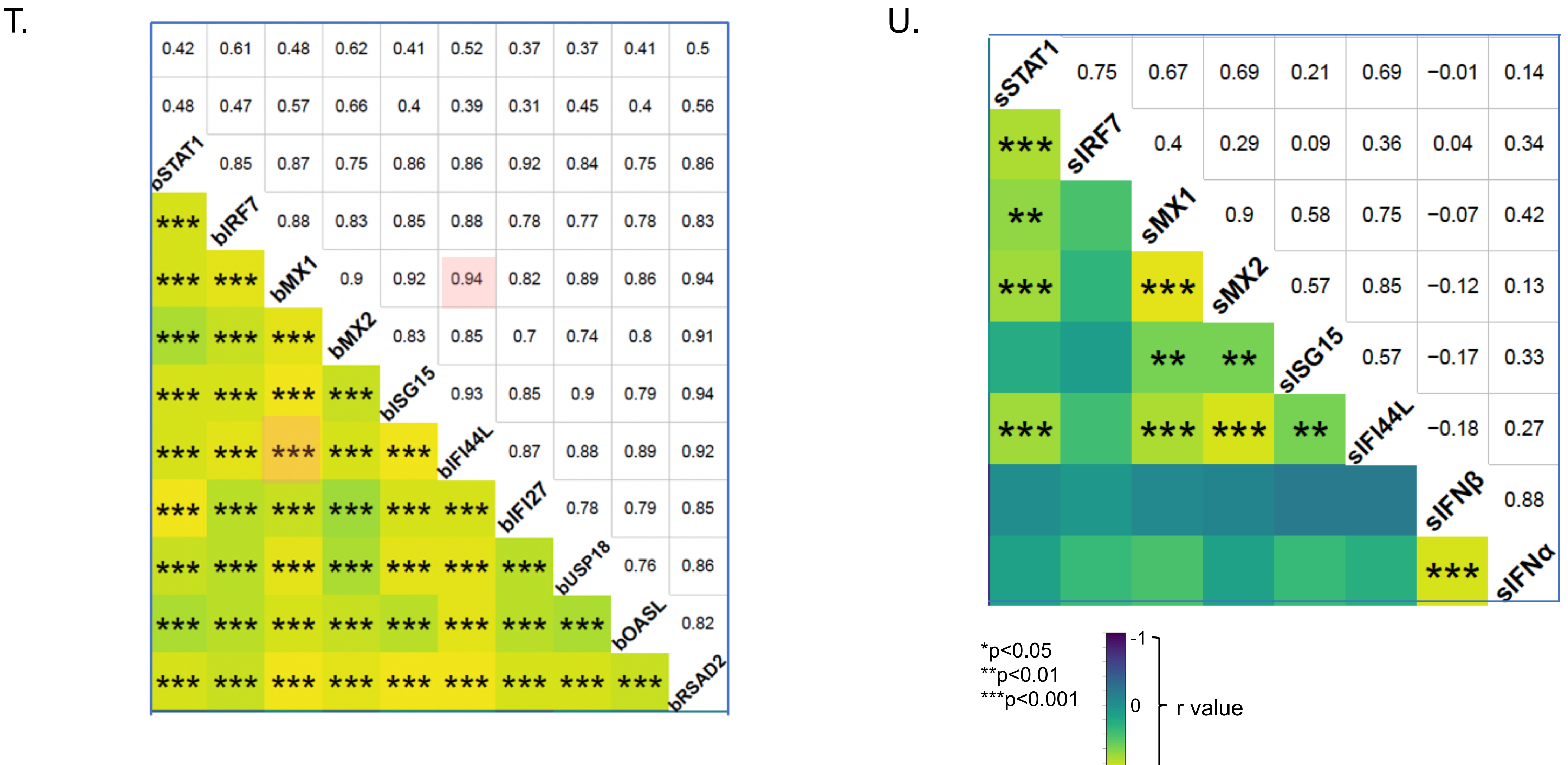


**Supplemental Figure 4T-U Correlation matrix for ISGs,** at baseline/unstimulated within an hour of sampling (A) and after 18 hours of stimulation with type I IFN. Numerical values refer to r value (Pearson's correlation) and ** to p value. Color intensity related to strength pf correlation between the two parameters eg basal MX1 expression is strongly correlation with basal IFI44L expression in freshly isolated monocytes from IPF patients; r=0.94 and p<0.001 (pink boxes in (A). 's' refers to stimulated; 'b' to basal gene expression by qPCR (values shown in Fig. 4)

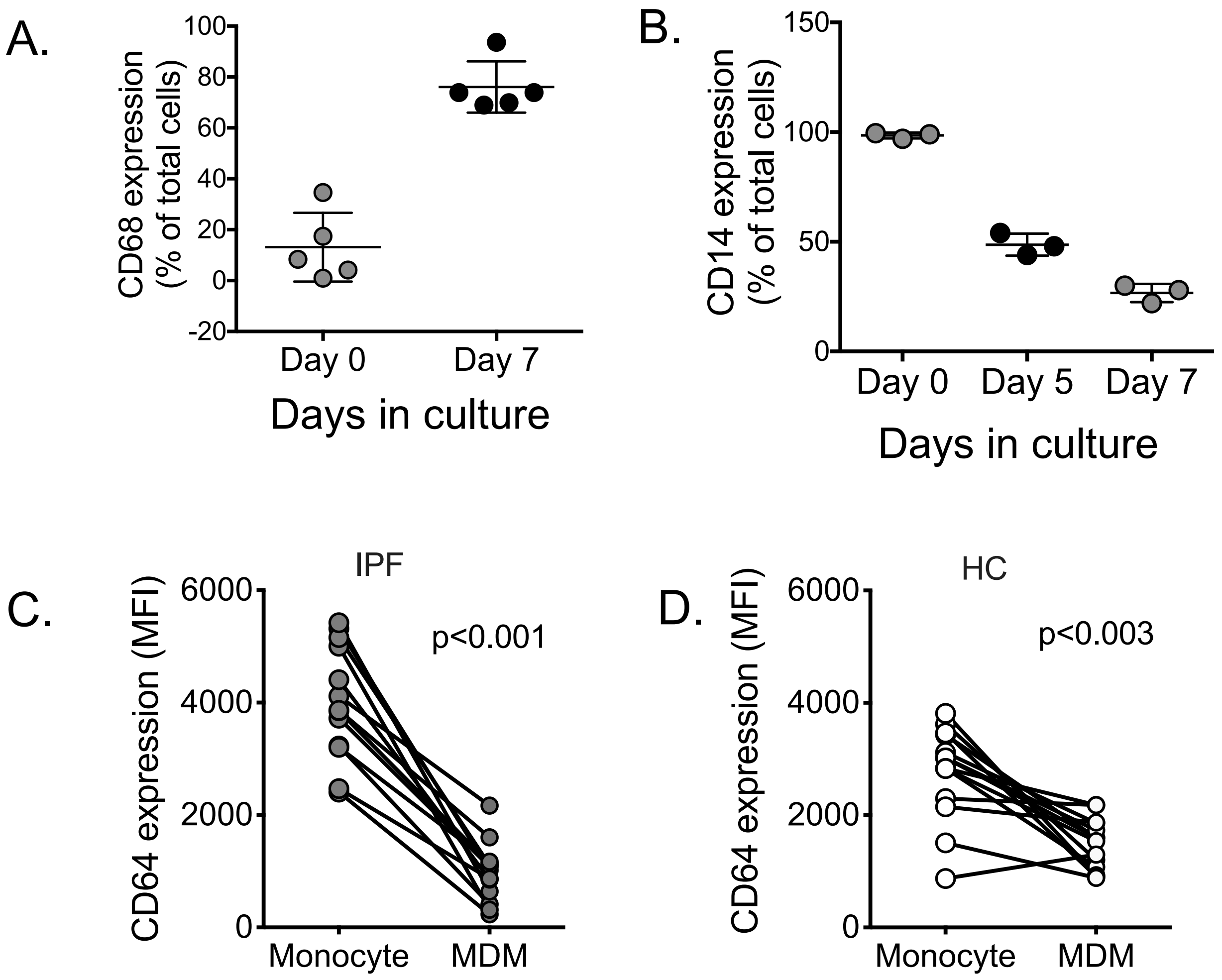


**Supplemental Figure 5. Changes in CD14, CD68 and CD64 on monocytes during maturation to MDMs.** FACS results from 7 day monocyte culture in 10% autologous serum and after a single CSF-1 dose for three healthy controls. (C-D) Changes in CD64 expression (by MFI) on during 7 days culture for IPF monocytes to macrophages (C) and for HC (D).

C.

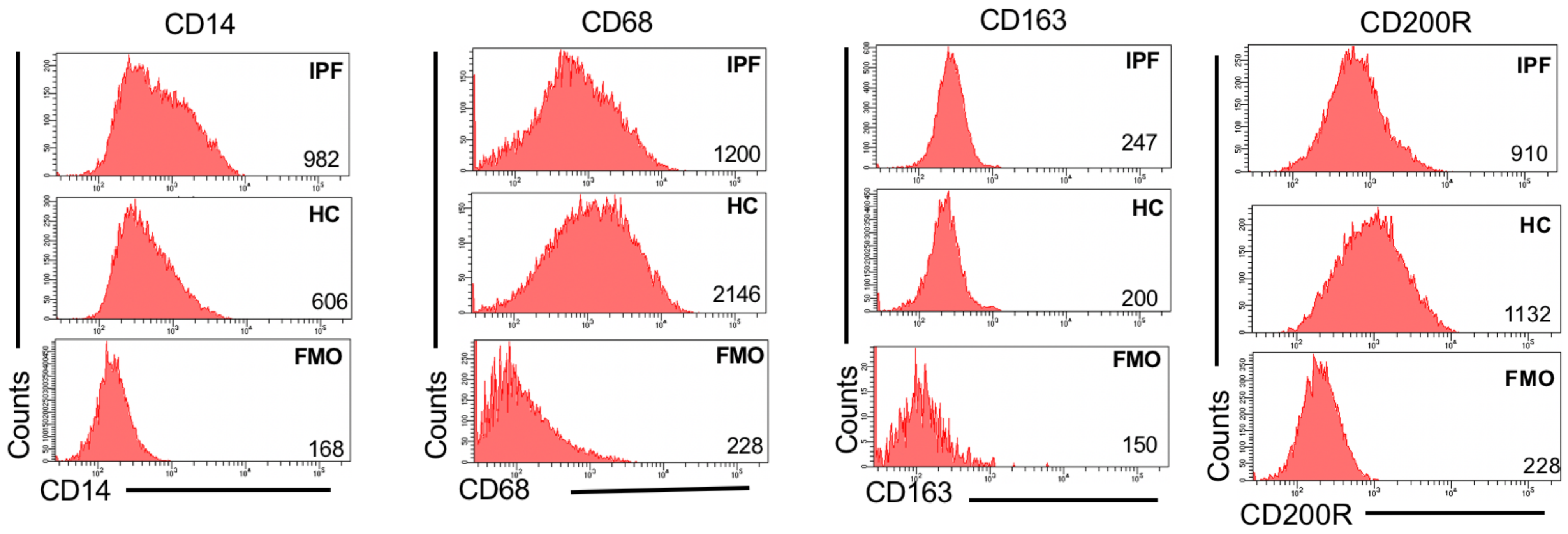


MDM after 7 days culture

**Supplemental Figure 5C Representative histogram of CD14, CD68, CD163 and CD200R expression on MDMs;** by flow cytometry after 7 day culture in 1% autologous serum and after a single CSF-1 dose

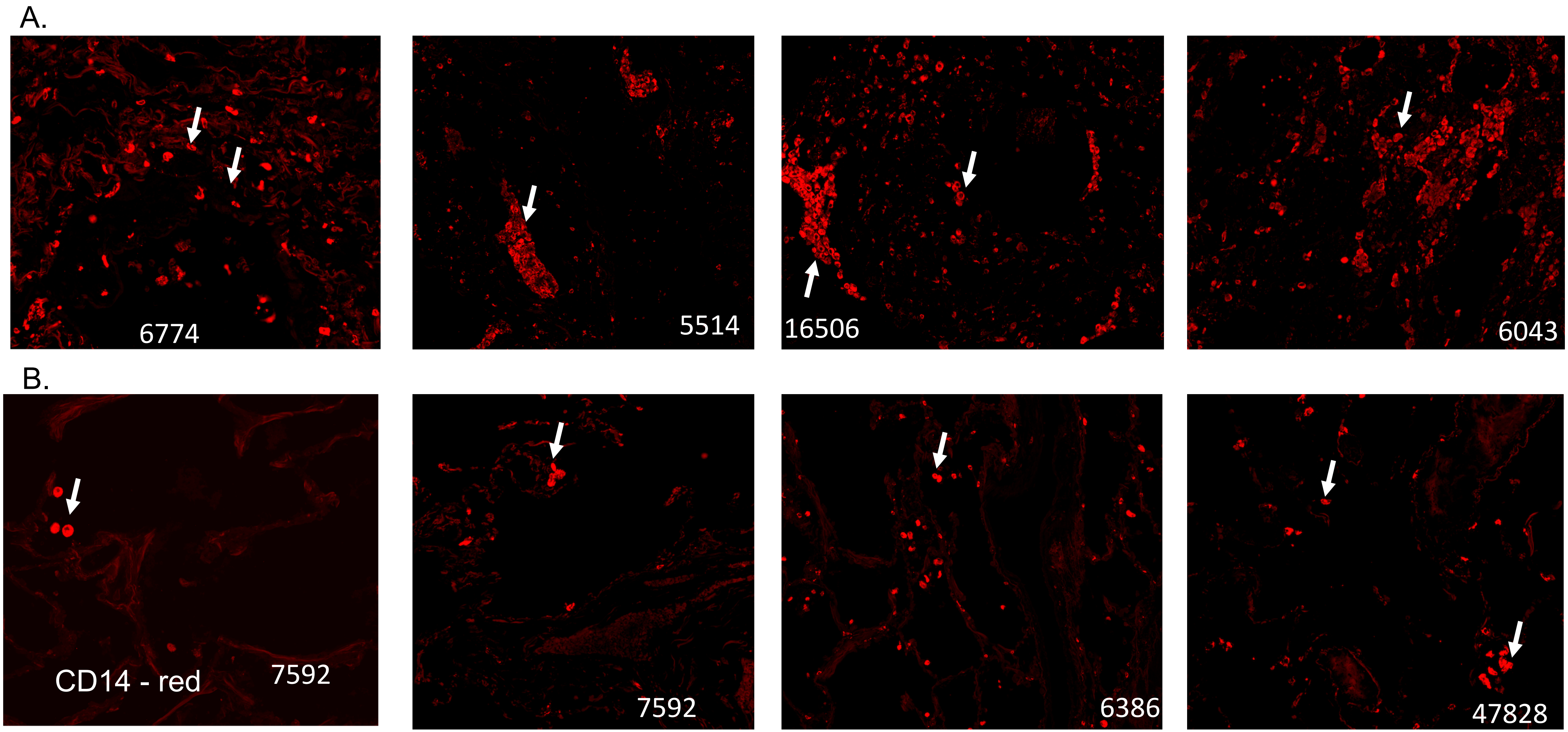


**Supplemental Figure 6A-B**. **CD14 expressing cells in IPF lungs.** CD14 immunofluorescent staining of paraffin embedded lung sections from IPF and control lungs. Red – CD14. Top panel (A) – 4 different lung samples from IPF patients. Bottom panel (B) non-IPF control lungs from 4 patients. Numbers refer to anonymous identifiers. Arrows indicate CD14 expressing cells which can be monocytes or monocytes differentiating to macrophages

C.

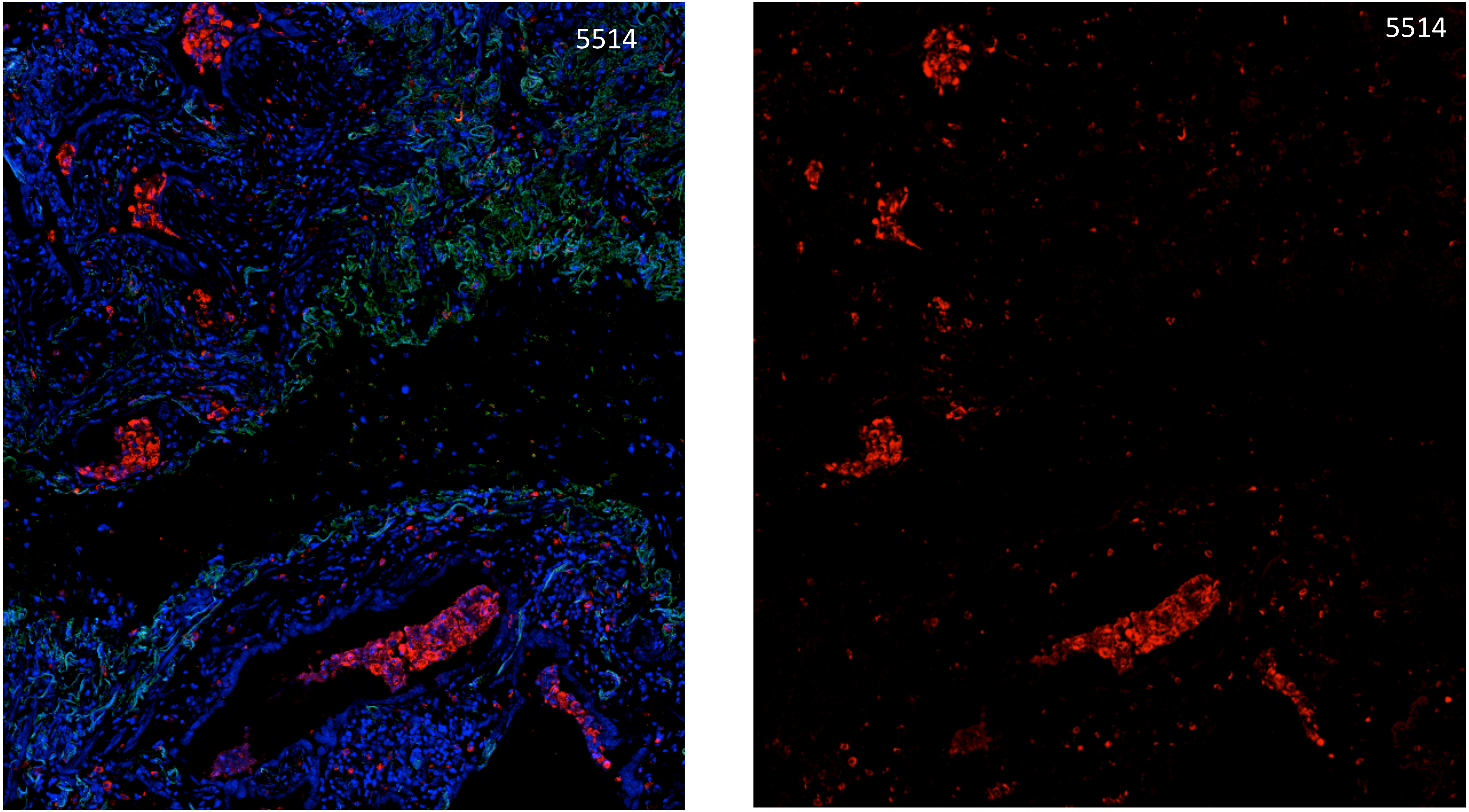


**Supplemental Figure 6C. CD14 expressing cells in IPF lungs.** CD14 expression (red) in IPF lung from patient 5514 with blue showing DAPI, and green [illegible]

D.

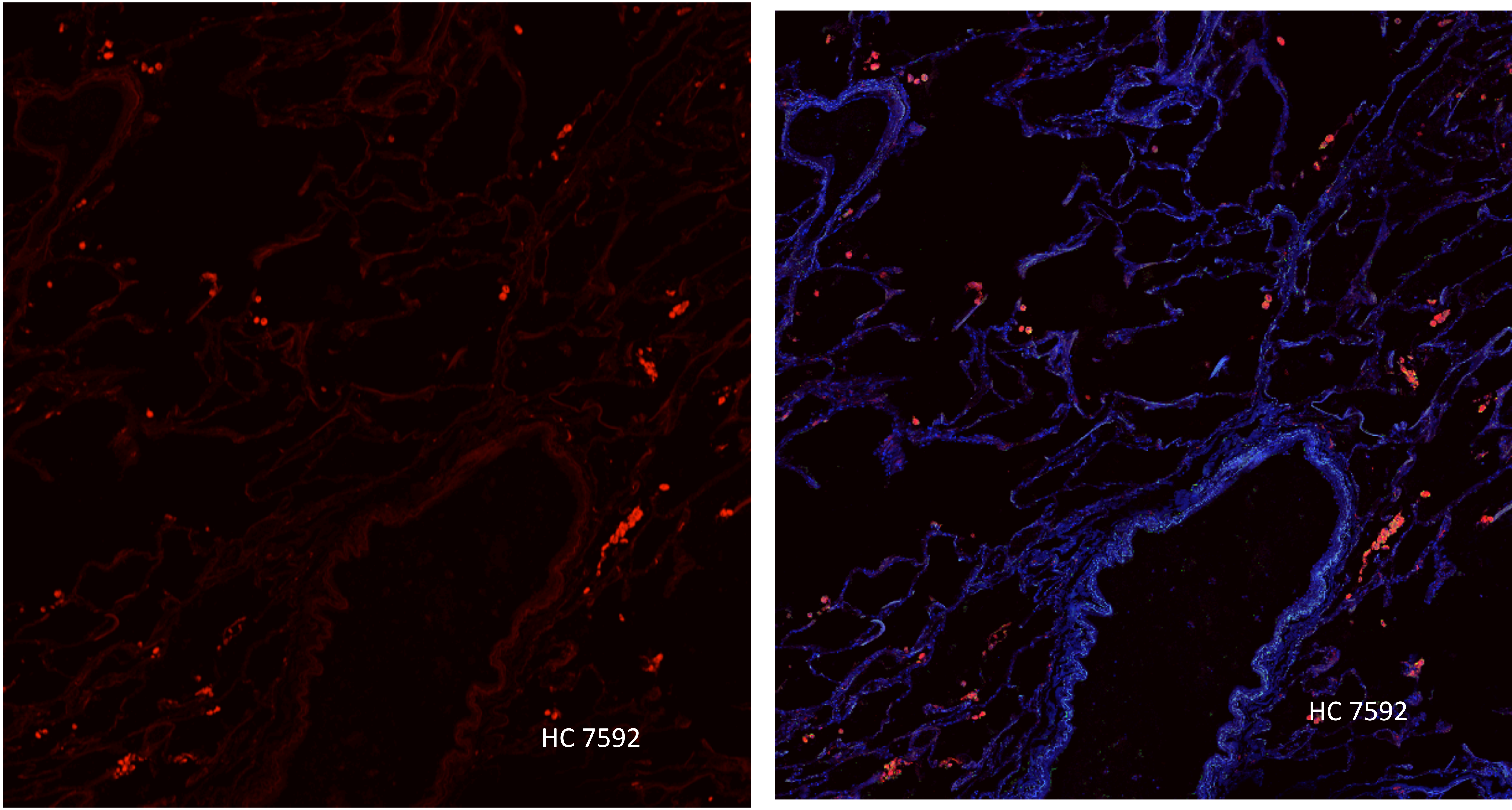

| IPF cohort | IPF | Controls |
|---|---|---|
| n | 28<br>(37 samples) | 28<br>(28 samples) |
| Male | 23 | 19 |
| Mean age (range) | 72.9y<br>(57-87) | 66y<br>(44-81) |
| On Pirfenidone | 14 | N/A |
| Mean FVC<br>(range) | 71<br>(48-124) | N/A |
| Mean TLCO<br>(range) | 48<br>(18-90) | N/A |
| Mean FEV1<br>(range) | 73<br>(40-107) | N/A |
| Mean CPI<br>(range) | 47<br>(15-6) | N/A |

Supplemental table 1

Demographic data for patient cohorts

| AEIPF cohort | AEIPF |
|---|---|
| Number of samples | 10 |
| Male (%) | 8 (80) |
| Mean age (range) | 71y<br>(52-80) |
| Definite UIP | 5 |
| Probable UIP | 5 |
| On Pirfenidone | 6 |
| On Prednisolone | 10 |

Supplemental table 2A

| Sample code | Time since diagnosis (years) | Length of symptoms prior to sampling (weeks) | No. of previous admissions for exacerbations | No. of days on Prednisolone prior to sampling | Pirfenidone (P)/ Nintedanib (N) treatment | CT findings | Outcome |
|---|---|---|---|---|---|---|---|
| 01BWAE | 5 | 6 | 3 | >10 | Yes (P) | New patchy GG | Home for palliation<br>RIP 19/06/2015 |
| 02AEAE | 2.8 | 3 | 1 | >10 | Yes (P) | Widespread GG | RIP 7/3/2015 |
| 03NKAE | 2.5 | 2 | 1 | >10 | Yes (P) | New GG<br>Progressive HC | Home<br>RIP 11/3/2016 |
| 04PFAE | 5 | 5 | 0 | 14 | No | Widespread GG<br>Patchy consolidation | Hospital RIP 24/4/15 |
| 05JFAE | 4.5 | 3 | 0 | 5 | Yes (N) | Widespread GG<br>Progressive HC | Home<br>RIP 19/8/2015 |
| 06PBAE | 0 | 3 | 0 | 2 | No | Patchy GG<br>Progressive HC | Home |
| 07CBAE | 4 | 2 | 0 | 6 | No | Airspace opacification GG | Home for palliation<br>RIP 20/10/2015 |
| 08HHAE | 3.7 | 4 | 2 | >10 | No | Progressive HC Multifocal GG | Improvement and home<br>RIP 30-9-2016 |
| 09RSAE | 2.4 | 1 | 0 | 5 | Yes (P) | New GG | Initial improvement<br>RIP 3/11/2016 |
| 10JDAE | 3.5 | 4 | 2 | >10 | Yes (P) | Little GG<br>Small patch of consolidation | Home<br>Palliative care |

Supplemental Table 2B Clinical details of patients with AEIPF. GG- ground glass, HC- honeycomb, CT – computerized tomographic images, RIP- de

| Genes (M2-like/anti-inflammatory) | Protein transcript/role | Control Vs Stable IPF<br><br>P-value |
|---|---|---|
| *TGFß1* | Pleiotropic profibrotic cytokine (M2-like marker) | 0.056 |
| *IL10* | Immunomodulatory cytokine (M2-like marker) | 0.090 |
| *CD206* | Mannose scavenger receptor (M2-like marker) | Low/no expression |
| *CD200R1* | Glycoprotein receptor | 0.063 |
| *TGM2* | Tissue transglutaminase (M2 like marker) | 0.131 |
| *CD163* | Haemoglobin scavenger receptor (M2 like marker) | 0.152 |

| Genes (M1-like/pro-inflammatory) | Protein transcript/role | Control Vs Stable IPF<br><br>P-value |
|---|---|---|
| *TNFα* | Inflammatory cytokine (M1- like marker) | 0.097 |
| *IL6* | Inflammatory cytokine (M1 like marker) | 0.205 |
| *CXCL10/ IP-10* | Chemokine (M1 like marker) | 0.323 |
| *CCR2/ MCP1* | Receptor for CCL2 (MCP-1) | 0.536 |
| *IDO1* | Indoleamine 2,3-dioxygenase 1(M1- like marker) | 0.765 |
| *IL1R2* | Decoy receptor | 0.232 |

Supplemental Table 3 Results from qPCR of relevant genes  in monocytes from IPF vs age-matched healthy controls.

| Monocyte qPCR cohort | Controls | IPF |
|---|---|---|
| n | 8 | 7 |
| Mean age (range) | 67 (50-72) | 71 (57-82) |
| Male (%) | 7 | 7 |
| % on Pirfenidone | N/A | 2 |
| Monocyte purity | 98.6 (98.6-98.7) | 99.2 (97.8-99.8) |
| Mean RIN (Range) | 9.5 (8.9-10) | 10 (9.5-10) |

Supplemental Table 4

| Serum mediators cohort | IPF | Controls |
|---|---|---|
| n | 24 | 11 |
| Male | 18 | 5 |
| Mean age (range) | 74y (63-86) | 69y (49-86) |
| On antifibrotics | 8 | N/A |

Supplemental Table 5

Demographic data for patient cohorts for analysis of monocyte gene expression by qPCR and for measurement of serum mediators

A.

| Type I IFN baseline cohort | Controls | IPF |
| --- | --- | --- |
| n | 10 | 27 |
| Age<br>mean (range) | 67<br>(56-79) | 75<br>(54-84) |
| Male (n) | 6 | 23 |
| % on antifibrotics | NR | 15 |

B.

| Type I IFN (IFN-stimulated) cohort | Controls | IPF |
| --- | --- | --- |
| n | 10 | 20 |
| Age<br>mean (range) | 67<br>(56-79) | 76<br>(58-84) |
| Male (n) | 6 | 17 |
| % on antifibrotics | NR | 11 |

Supplemental Table 8. demographic for patient cohorts used for examination of type 1 IFN expression at baseline and after type I IFN stimulation

A

| MDM phenotyping cohort | IPF | Controls |
|---|---|---|
| n | 17 | 19 |
| Male (n) | 13 | 68 |
| Mean age (range) | 71.0 (62-87) | 64.9 (49-86) |
| % on anti-fibrotics | 5 | N/A |

B

| MDM groups | Phagocytosis | | ROS | | Neutrophil Efferocytosis | |
|---|---|---|---|---|---|---|
| | IPF | Controls | IPF | Controls | IPF | Controls |
| n | 19 | 12 | 9 | 7 | 12 | 15 |
| Male (n) | 13 | 58 | 9 | 6 | 11 | 10 |
| Mean age (range) | 75 (63-87) | 66.1 (44-80) | 73 (58-82) | 67 (57-73) | 77 (66-85) | 65.6 (57-75) |
| On antifibrotic (n) | 7 | NA | 5 | NA | 3 | N/A |

Supplemental Table 9. demographic for patient cohorts used for monocyte-derived macrophage (MDM) experiments

| Reference | Age (years) | MDT diagnosis | Smoking | Treatment at point of sampling | FEV1 % | FVC% | DLCO% |
|---|---|---|---|---|---|---|---|
| HC1_92 | 66 | T1aN0M0 adenoca | Ex smoker > 5 years ago | none | 80 | 96 | 62 |
| HC2_86 | 82 | T1bN0M0 adenoca | Never smoker | none | | | |
| HC3_28 | 70 | T1bN0M0 adenoca | Never smoker | none | 120 | 124 | 108 |
| HC4_78 | 78 | T1aN0M0 adenoca | Ex smoker > 5 years ago | none | 64 | 97 | 84 |
| IPF1_43 | 37 | IPF | Ex smoker > 5 years ago | N-acetyl cysteine | 63 | 60 | 48 |
| IPF2_74 | 63 | IPF | Ex smoker > 5 years ago | none | 68 | 69 | 65 |
| IPF3_06 | 62 | IPF | Ex smoker > 5 years ago | none | 78 | 73 | 54 |
| IPF4_14 | 76 | IPF | Ex smoker > 5 years ago | none | 109 | 98 | 52 |

Supplemental Table 10. Demographic for patient cohort used for lung sampling. TNM - Tumour, Node, Metastasis staging system according to 7th Edition of TNM international staging (Goldstraw P 2009). Adenoca- adenocarcinoma
